# Exploring endocytic compartment morphology with systematic genetics and single cell image analysis

**DOI:** 10.1101/724989

**Authors:** Mojca Mattiazzi Usaj, Nil Sahin, Helena Friesen, Carles Pons, Matej Usaj, Myra Paz Masinas, Ermira Shuteriqi, Aleksei Shkurin, Patrick Aloy, Quaid Morris, Charles Boone, Brenda J. Andrews

## Abstract

Endocytosis is a conserved process that mediates the internalization of nutrients and plasma membrane components, including receptors, for sorting to endosomes and the vacuole (lysosome). We combined systematic yeast genetics, high-content screening, and neural network-based image analysis of single cells to screen for genes that influence the morphology of four main endocytic compartments: coat proteins, actin patches, late endosome, and vacuole. This unbiased approach identified 17 mutant phenotypes and ∼1600 genes whose perturbation affected at least one of the four compartments. Numerous mutants were associated with multiple phenotypes, indicating that morphological pleiotropy is often seen within the endocytic pathway. Morphological profiles based on the 17 aberrant phenotypes were highly correlated for functionally related genes, enabling prediction of gene function. Incomplete penetrance was prevalent, and single-cell analysis enabled exploration of the mechanisms underlying cellular heterogeneity, which include replicative age, organelle inheritance, and stress response.

---

Endocytosis is a highly conserved bioprocess that plays a central role in eukaryotic cell biology, mediating the internalization of receptors, nutrients, and other molecules, controlling the lipid and protein composition of the plasma membrane and the coupling of different intracellular compartments^1^. Endocytosis initiates with vesicle formation at specific sites at the plasma membrane. In yeast, proteins that are required for vesicle formation have been grouped into five functional modules based on their dynamic behaviour: the early proteins, the coat module, the WASp/myosin module, the actin module and the scission module^2^. Coat proteins perform several roles, including cargo uptake and regulation of actin dynamics, functioning as adaptors to link cargo, coat, plasma membrane and actin network components. Actin patches represent a later stage in internalization; their appearance coincides with the membrane invagination and coat internalization step^3^. The modular design of yeast endocytic vesicle formation is largely recapitulated in mammalian cells^4^. After cargo uptake, endocytic vesicles fuse with early endosomes, allowing cargo to be recycled to the plasma membrane, or targeted through more mature endosomes and multivesicular bodies (MVBs) for vacuolar (lysosomal) degradation. The endocytic intracellular trafficking pathway impinges on a number of cellular physiological processes and is often associated with the pathology of human diseases, including atherosclerosis, some cancers, and Alzheimer’s disease^5, 6^.

Large-scale genetic screens have been combined with cell biological analysis to explore different aspects of endocytic trafficking in yeast and higher eukaryotes. These studies have defined several core components and regulators of the endocytic pathway^7–10^. However, most cell biological approaches applied thus far use population-level measurements as a phenotypic read-out, which precludes quantitative analysis of cell-to-cell heterogeneity^11^. Here we explore the yeast endocytic pathway using systematic genetic analysis combined with high-content screening. We examined 5292 unique yeast genes for roles in endocytic compartment morphology, applying live-cell fluorescence microscopy and neural network-based, single-cell image analysis. In total, we identified ∼1600 genes whose perturbation affects at least one endocytic compartment, revealing both new biology and insights into mechanisms underlying cellular heterogeneity. The experimental and computational pipeline developed here can be generalized to other unrelated compartments, pathways or phenotypes, which will allow us to expand our knowledge on the inner workings of a cell. Importantly, the computational analysis framework we developed is also species independent, and we provide the tools for its implementation.

## RESULTS

### Combined experimental-computational pipeline for quantitative single-cell assessment of mutant phenotypes of endocytic compartments

To enable a quantitative analysis of the morphology of the main endocytic compartments, we developed a high-throughput (HTP) image-based pipeline coupled to single-cell image analysis (Figure 1a). We constructed a series of query strains with a fluorescent protein (FP) at the C-terminus of four endogenous yeast proteins, each serving as a marker for a unique endocytic compartment. We focused on: [1] *SLA1*, encoding an endocytic adaptor protein, marking the coat complex associated with early endocytic sites at the plasma membrane; [2] *SAC6*, encoding yeast fimbrin, marking actin patches that are also required for early endocytosis events; [3] a late endosomal marker, *SNF7*, encoding a subunit of the ESCRT-III complex involved in the sorting of transmembrane proteins into the multivesicular body (MVB) pathway; and [4] a marker for the vacuolar membrane, *VPH1*, encoding subunit ‘a’ of the vacuolar ATPase (V-ATPase) V_O_ domain (Figure 1b).

**Figure 1:**
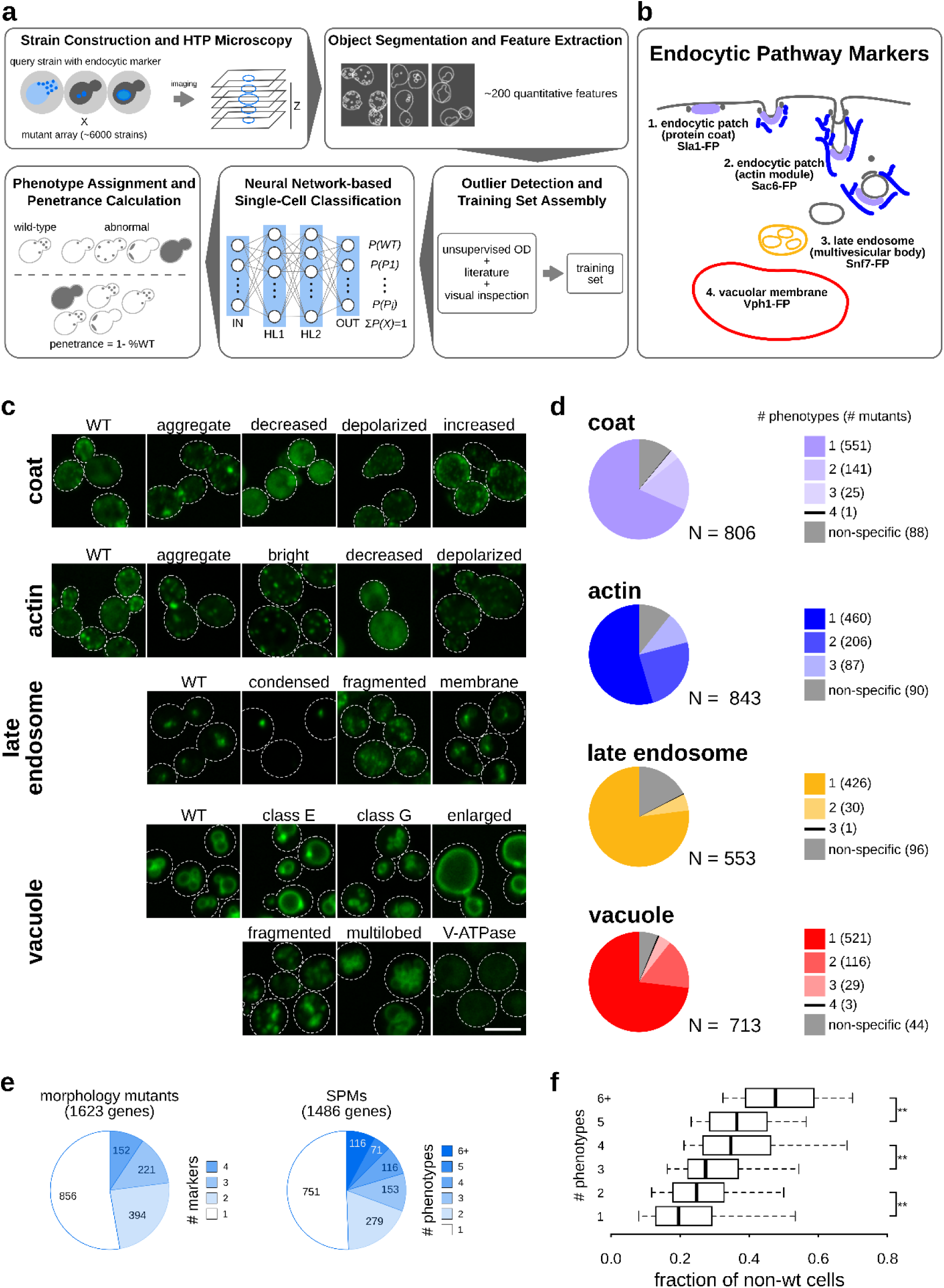
Twenty-one subcellular endocytic phenotypes identified using computational analysis of single cell images. (see also Supplementary Figure 1, Supplementary Figure 2, Supplementary Table 1, Supplementary Table 2). (a) Diagram of the experimental and computational workflow. Yeast mutant arrays harbouring fluorescently-tagged proteins marking specific endocytic compartments were constructed using the synthetic genetic array (SGA) method and imaged using automated high-throughput microscopy. Image and data pre-processing steps included object segmentation and feature extraction, low-quality object clean-up and data standardization. Positive controls and classification training sets were used to train a fully connected 2-hidden-layer neural network (2NN), allowing assignment of phenotypes at the single-cell level and calculation of penetrance. (b) Illustration of endocytosis process and compartment markers. The four endocytic compartment markers used in this study are indicated: Sla1 as a marker of the protein coat component of the endocytic patch (light purple); Sac6 as a marker of the actin component of the endocytic patch (blue); Snf7 as a marker of the late endosome (orange); Vph1 as a marker of the vacuolar membrane (red). The colours chosen for each marker are used throughout this study. FP: fluorescent protein. (c) Example micrographs of yeast cells for each of the 21 subcellular endocytic phenotypes identified in this study. The relevant markers are listed to the left of the micrographs. Scale bar: 5 µm. (d) Pie charts showing the proportion of specific phenotype mutants (SPMs) that have one or more distinct aberrant phenotypes, and non-phenotype-specific mutants for each of the compartments screened. (e) Pie charts showing the proportion of mutant strains that are morphology mutants for one or more markers (left) and specific phenotype mutants (SPMs) that cause one or more aberrant morphological phenotypes (right). The number of mutants in each category is listed within each section. (f) Box plot illustrating the distribution of the fraction of non-wild-type (WT) cells for specific phenotype mutants grouped by the number of phenotypes they cause. ** denotes a significant difference between two groups (p-value < 0.01; significance was determined using analysis of variance (ANOVA) with a post-hoc Bonferroni test). Whiskers extend to the 5^th^ and 95^th^ percentile.

We introduced each marker into both the yeast deletion collection^12^, and the collection of temperature-sensitive (TS) mutants of essential genes^13, 14^, using the synthetic genetic array (SGA) approach^15^. We acquired live cell images of log phase cultures with an automated HTP microscope. CellProfiler^16^ was used to identify individual cells and subcellular compartments, and extract quantitative features describing these segmented compartments. The final dataset included quantitative data for ∼16.3 million cells from 5627 mutant strains (5292 unique ORFs or ∼90% of yeast genes), with an average of 640 cells for each mutant strain.

To find mutants with abnormal compartment morphology, we used an automated method to identify “outlier” cells with non-wild-type morphology (see Methods) and then visually inspected strains with many outlier cells to identify common patterns of morphological defects that we used to define phenotypes. From these strains, and a set of positive control strains for mutants with known endocytic defects (Supplementary Table 1), we defined 21 endocytic phenotypes: 4 wild-type, one per compartment, and 17 showing aberrant morphology (Figure 1c).

We then labelled a representative set of cells displaying these 21 phenotypes using a custom-made, single-cell labelling tool; this “training set” was used to train a neural network to automatically classify other cells. To confirm that the CellProfiler features derived from the cell images were sufficient to distinguish the different mutant phenotypes, we performed non-linear dimensionality reduction using t-SNE^17^ on the training set feature vectors, and confirmed that cells with different phenotypes occupied distinct regions of the 2D reduced feature space (Supplementary Figure 1a). We then used the labelled dataset to train a 2-hidden-layer fully connected neural network (2NN) for each of the endocytic markers. For each single cell, the marker-associated 2NN estimated the probability of each phenotype and we assigned each cell the phenotype with the highest probability. We used CellProfiler features instead of those learned using a convolutional neural network (CNN) because, unlike recent studies^18–21^, the CNN performed poorly on our relatively small training set (data not shown). The average classification accuracy on held-out data across all markers and phenotypes was 88.4%, and 18 of the 21 phenotypes had an average classification accuracy >80% (Supplementary Figure 1b-c, Supplementary Table 1, see Methods).

Statistical analyses validated the quality of our pipeline, confirming reproducibility and accuracy of the single cell phenotypic classifications (see Methods; Supplementary Figure 1d-h). Applying our 2NN to the entire dataset allowed us to accurately detect even a small fraction of aberrant cells, enabling quantification of the variety and penetrance of mutant phenotypes associated with a given mutation (see below).

### Hundreds of yeast genes affect endocytic compartment morphology

To capture the spectrum of phenotypes associated with each mutant strain, we determined the fraction of cells in a mutant strain population that displayed each of the 21 phenotypes using our classifiers described above (Figure 1c). We called a strain a specific phenotype mutant (SPM) if the fraction of cells assigned an aberrant phenotype was significantly greater than that assigned the same phenotype in a control wild-type strain population (see Methods). In total, we identified 1486 mutants as SPMs (Figure 1d), with many mutants classified as SPMs for more than one phenotype. We defined a subset of 363 mutants as stringent SPMs, as they had a relatively larger fraction of cells with a specific defect (see Methods). We also identified a small set of non-phenotype-specific mutants (137 unique genes; Figure 1d) which showed a significant increase in the total percentage of the cell population displaying an aberrant phenotype for a given compartment, even if none of the individual phenotype fractions were high enough for a given strain to be classified as an SPM.

In total, we identified 1623 yeast genes (∼30% of screened ORFs) that affect the morphology of one or more endocytic compartments (referred to as morphology mutants; Supplementary Table 2). Thus, yeast endocytosis is remarkably sensitive to single gene perturbation, consistent with previous siRNA screens in mammalian cells^9^.

For each marker, some of the morphology mutants showed multiple phenotypes (Figure 1d). Overall, approximately half of the 1623 morphology mutants showed aberrant phenotypes with more than one of the four markers screened, and approximately half of the SPMs displayed more than one of the 17 aberrant phenotypes (Figure 1e), indicating that morphological pleiotropy, which we define as occurring when a mutant has two or more aberrant morphological phenotypes, is prevalent within the endocytic pathway and that numerous genes impinge on multiple stages of endocytosis. The most pleiotropic mutants (those causing six or more specific phenotypes; 116 SPM genes) were involved in vesicle organization, exocytosis, protein lipidation, and membrane fusion. Genes associated with multiple morphological outcomes tended to affect a larger fraction of the cell population (Figure 1f). Morphology mutants were also enriched for TS alleles of essential genes (Supplementary Figure 2a-b) and the fraction of essential gene mutants increased with the number of morphological phenotypes (Supplementary Figure 2c). However, morphological pleiotropy was not confined to essential genes. For example, mutants of both the essential exocyst complex and the nonessential ESCRT complexes led to phenotypic defects spanning the early and late endocytic compartments (Supplementary Table 2).

Genes annotated with roles in a wide range of functions appear to impinge on the endocytic pathway. Only 286 (∼18%) of the identified mutant genes were annotated to GO Slim biological process terms associated with endocytosis and the endomembrane system (Supplementary Table 2). Similarly, while morphology mutants were enriched for genes conserved between yeast and human (∼40% of conserved morphology mutants compared to ∼26% on the array, p-value < 0.0001; Supplementary Figure 2d), this enrichment was not due to known endocytosis machinery components (Supplementary Figure 2e), but included genes involved in a range of bioprocesses, such as DNA replication and repair, transcription and splicing.

### Automated image analysis identifies the spectrum of possible endocytic compartment morphologies

Of the 17 aberrant morphological phenotypes associated with the four endocytic markers, 15 correspond to previously described phenotypes (Supplementary Table 1). The unsupervised outlier detection analysis identified two novel phenotypic groups: mislocalization of the late endosomal marker to the vacuolar membrane (“late endosome: membrane” in Figure 1c), and a previously unappreciated vacuolar mutant phenotype characterized by small vacuoles and increased cytosolic localization of the vacuolar marker, Vph1. Because most of the SPMs in this class were genes involved in various aspects of Golgi vesicle transport, we refer to this vacuolar morphology as the ‘class G’ phenotype. We confirmed that the class G was a distinct phenotype, and not an intermediate stage of one of the known vacuolar phenotypes, by imaging in a 24h time-course at 37°C (Supplementary Figure 3a). Since Golgi vesicle transport affects trafficking pathways to the vacuole, the class G phenotype could be a consequence of abnormal vacuolar membrane composition that leads to defects in vacuole formation or membrane fusion and fission.

Comparisons to a panel of gene attributes (Supplementary Figure 3b, Supplementary Table 3) revealed that morphology mutants in all four compartments were enriched for the same set of features: high conservation across different species, ample genetic interactions (GIs) and protein-protein interactions (PPIs), pleiotropy and multifunctionality, enrichment for fitness defects, and tendency to act as phenotypic capacitors.

Mutants with aberrant phenotypes were often enriched in multiple bioprocesses, both closely related and apparently unrelated to the compartment associated with the aberrant phenotype, suggesting that multiple mechanisms can lead to a particular phenotype (Figure 2, Supplementary Table 3). Stringent SPMs were enriched for more specific protein complexes and biological pathways, which may be suggestive of the mechanisms underlying their aberrant morphological phenotypes (Supplementary Table 3). Phenotypes that occur in a relatively high fraction of the population in wild-type strains, such as depolarized patches or multilobed vacuoles (Figure 2), may result from a general cellular response to different stress conditions (environmental or genetic). These phenotypes tend to be associated with a larger number of SPMs (Figure 2), whereas phenotypes with few SPMs more likely indicate responses associated with a specialized pathway.

**Figure 2:**
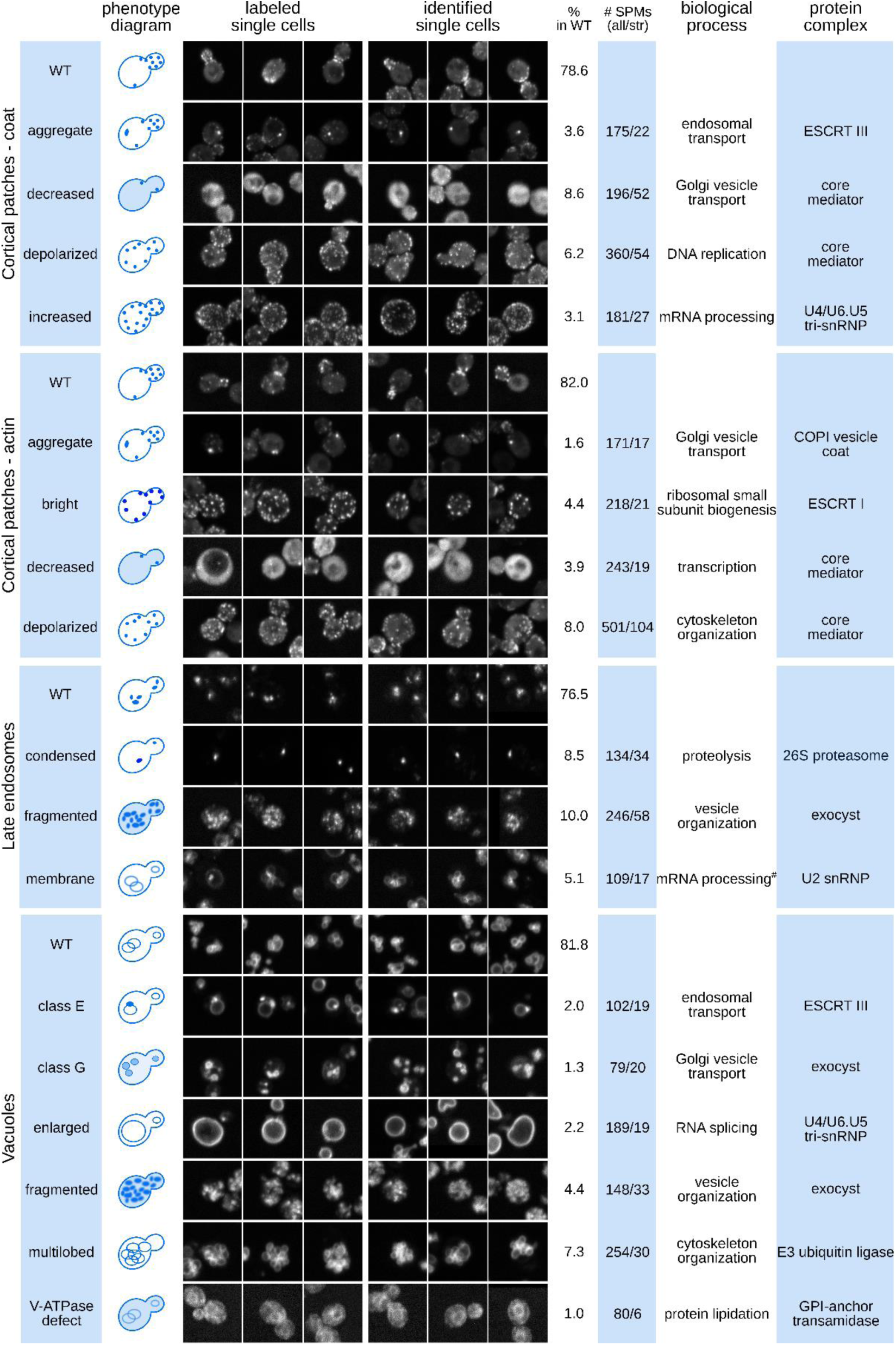
The spectrum of endocytic compartment morphologies: properties of 17 mutant phenotypes. (see also Supplementary Figure 3, Supplementary Table 3). Representative images of wild-type and mutant cells organized by marker and phenotype (labels on the left of each panel). For each phenotype, three cells labelled for the training set (labelled single cells) and cells identified by the 2NN classifier (identified single cells) are shown. The table to the right of the images shows (from left to right): [1] the occurrence of each phenotype in a wild-type population (% in WT); [2] the number of specific phenotype mutants (all) and stringent specific phenotype mutants (str) for each of the 17 mutant morphologies; [3] the most significantly enriched GO Slim biological process and; [4] the most significantly enriched protein complex. # denotes term below statistical significance.

The comparison of SPMs for several markers allowed us to search for new connections both within and between the endocytic compartments. We found that different morphology defects can be enriched among genes with roles in the same bioprocesses (Supplementary Table 3), possibly reflecting a common biological mechanism. We evaluated whether pairs of phenotypes shared more stringent SPMs than expected by chance (Supplementary Table 5). Of the 136 possible phenotype pairs, 36 pairs shared a significantly (p-value < 0.05, FDR < 0.2) overlapping set of causative gene mutations, and for 15 of these pairs the overlapping set was enriched in specific protein complexes (Figure 3a, Supplementary Table 5). This analysis identified a core set of 13 protein complexes that affect endocytic compartment morphology at multiple levels (Figure 3b). Some of the related endocytic morphology defects are likely sequential, while others may stem from independent events. For example, mutations in genes encoding components of the ESCRT complexes caused three connected phenotypes: coat aggregates, condensed late endosomes, and class E vacuoles. Defects in ESCRT complex assembly and MVB formation lead to accumulation of cargo at the late endosome - all three phenotypes therefore mark an exaggerated prevacuolar endosome-like compartment^22^. In contrast, mutation of genes encoding general transcriptional regulators such as TFIIH and the core mediator caused pleiotropic endocytic phenotypes which may reflect a series of independent defects in transcription.

**Figure 3:**
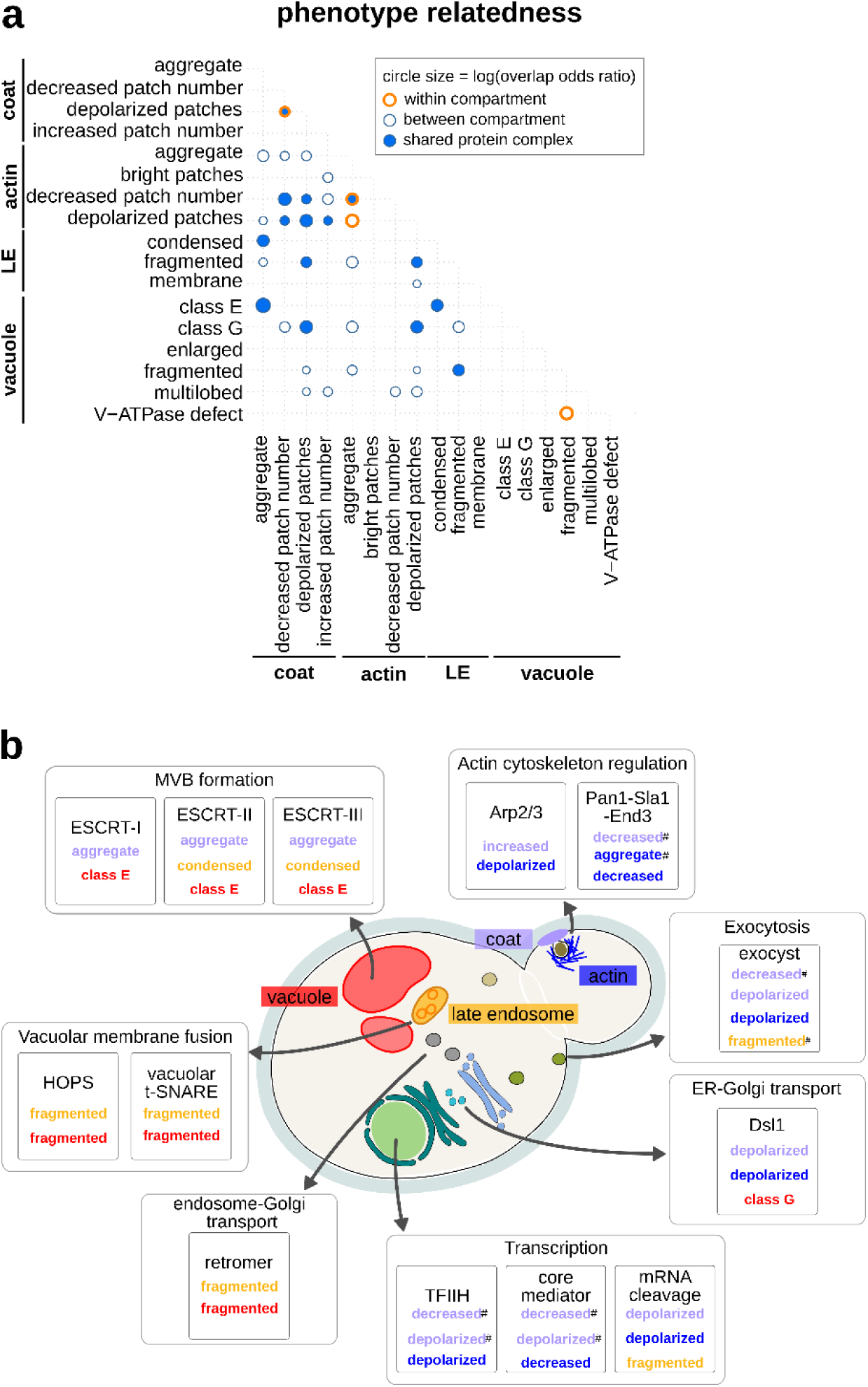
Analysis of the common morphology mutants of endocytic compartment phenotypes and the relationship to known protein complexes. (see also Supplementary Table 5). (a) Matrix showing significant overlap of stringent specific phenotype mutants (p-value < 0.05; significance was determined using Fisher’s exact tests). Circle size corresponds to the log value of the overlap odds ratio. Orange circles denote same-compartment phenotype pairs. Dark blue fill colour indicates phenotype pairs with at least one enriched protein complex in the overlapping set. LE: late endosome. (b) Diagram illustrating co-occurrence of endocytic morphology phenotypes associated with protein complex perturbation. Shown are significant protein complexes from (a) with biological processes and linked phenotype pairs. # denotes a phenotype pair without significant enrichment. Phenotype names are color-coded by endocytic marker, using the colour key described in Figure 1 and indicated on the yeast cell diagram.

To explore the extent to which the 17 aberrant endocytic compartment morphologies translate into a defect in endocytic internalization, we compared our list of SPMs with the published results of a quantitative assay for endocytic recycling of the non-essential gene-deletion collection, based on a GFP-Snc1-Suc2 chimeric protein^8^. All sets of SPMs derived from the 17 aberrant phenotypes were associated with a decrease in endocytic internalization (p-value < 0.01; Supplementary Figure 3c, Supplementary Table 6), with the exception of SPMs for the vacuolar class G phenotype. This observation may be due to a lack of power, because the class G SPMs were mostly essential genes (58/79 SPMs, including all of the stringent SPMs, Supplementary Figure 2b). We next tested each phenotype class to determine whether the mutants with a more penetrant version of the phenotype were more likely to have an endocytic internalization defect. We compared the defect levels of stringent SPMs to non-stringent SPMs, and found a significant increase in defects for four phenotypes: decreased number of actin patches, coat aggregate, condensed late endosome and class E vacuole (Supplementary Figure 3c, Supplementary Table 6). These phenotypes are likely directly linked to an endocytic internalization defect. The internalization defects of the stringent SPMs for actin patches were the highest of the four compartments (Supplementary Table 2, Supplementary Table 6). The actin module is the driving force in endocytic internalization and studies have previously shown that mutants with a reduced number of actin structures have defective endocytosis^3^. The remaining three phenotypes linked with internalization defects were those associated with defects in ESCRT complex and MVB formation.

### Subcellular morphology information and phenotype profiles support prediction of gene function

For virtually all 17 aberrant morphological phenotypes, we found several genes that had not been previously linked to the assessed morphological defects, including ∼130 morphology mutants corresponding to largely uncharacterized genes. For example, *YDL176W* caused a decrease in the number of actin patches and concomitant increase in the number of coat patches when mutated. This suggests a defect in actin patch assembly that causes a delay in patch internalization and accumulation of upstream components. Indeed, a *ydl176w*Δ mutant harbouring Sla1-GFP and Sac6-tdTomato markers exhibited a 55% increase in the lifetime of Sla1-GFP patches (p-value < 0.0001) and a modest but significant increase in the lifetime of Sac6-tdTomato (7.6% increase, p-value *=* 0.0012) (Figure 4a). Moreover, the *YDL176W* deletion mutant has an endocytic internalization defect^8^, and *YDL176W* shows a strong negative GI with *SLA2*^13^, which encodes an adapter protein that links actin to clathrin and endocytosis. We thus named the *YDL176W* open reading frame *IPF1* for **i**nvolved in actin **p**atch **f**ormation.

**Figure 4:**
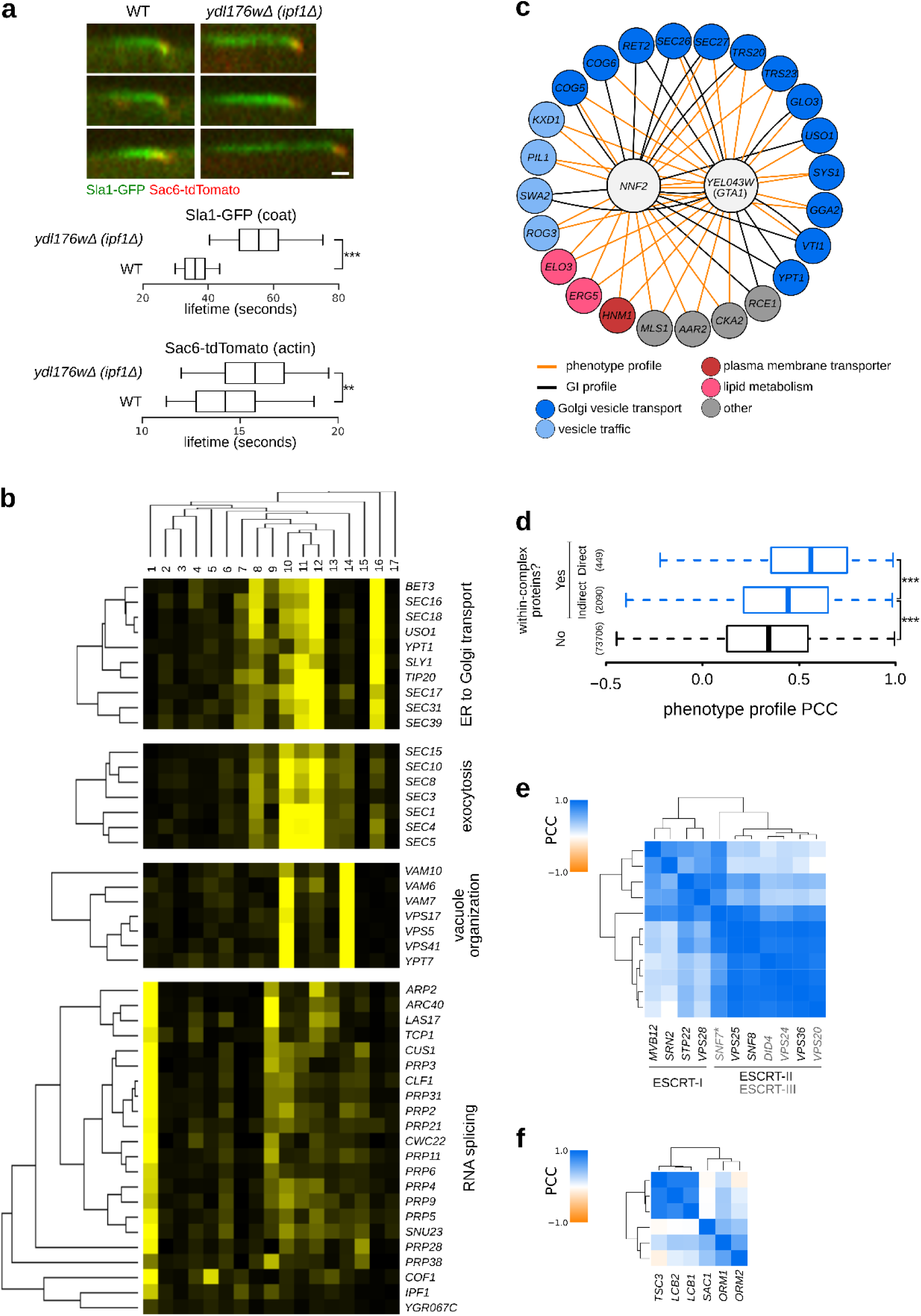
Predicting gene function from phenotype profiles. (see also Supplementary Figure 4). (a) Endocytic patch formation dynamics in the *ydl176w*Δ (*ipf1*Δ) strain. Patch dynamics were examined using time-lapse fluorescence microscopy of wild-type (WT) and *ipf1*Δ deletion strains carrying reporters for the coat (Sla1–GFP; green), and actin (Sac6-tdTomato; red) modules. Left: Representative kymographs for the WT and *ipf1*Δ strains. Right: Box plot illustrating the distribution of lifetimes of Sla1-GFP and Sac6-tdTomato patches. Significance was determined using unpaired t-tests. Scale bar: 10 seconds. (b) Examples of gene clusters obtained with hierarchical clustering of phenotype profiles composed of the 17 specific phenotype fractions. Phenotypes 1-17: [1] coat: increased patch number; [2] coat: aggregate; [3] vacuole: class E; [4] late endosome: condensed; [5] actin: bright patches; [6] late endosome: membrane; [7] actin: aggregate; [8] coat: decreased patch number; [9] actin: decreased patch number; [10] late endosome: fragmented; [11] coat: depolarized patches; [12] actin: depolarized patches; [13] vacuole: multilobed; [14] vacuole: fragmented; [15] vacuole: enlarged; [16] vacuole: class G; [17] vacuole: V-ATPase defect. (c) Interaction network of *NNF2* and *YER043W* (*GTA1*). Genes with phenotype profiles with a correlation > 0.7 and genetic interaction profiles with a correlation > 0.2, and at least two significant correlations to *NNF2* and/or *GTA1* were included in the network. (d) Analysis of phenotype profile similarity between mutants in genes encoding proteins in same or different protein complex structures. Box plot indicates distribution of PCCs between pairs of phenotype profiles for genes that encode protein pairs in direct contact in a protein complex experimental structure (Yes - Direct), code for protein pairs in the same protein complex structure but not in direct contact (Yes - Indirect), and code for protein pairs that do not belong to the same protein complex structure (No). The number of pairs evaluated in each set is shown on the x-axis. Significance was determined using one-sided Mann-Whitney U tests. ***p-value < 0.001. (e, f) Differentiation of functionally related protein complexes and protein complex organization using phenotype profiles. Heatmaps showing PCCs between components of the ESCRT complexes (e) and the SPOTS complex (f). A more intense blue colour indicates a higher PCC (scale bar at the top left of each heat map).

As we have shown, half of our SPMs affect multiple compartments and some lead to phenotypes that are present only in a small fraction of the population. To facilitate functional prediction for these genes, we used a multivariate approach that considers all the morphology phenotype classes. For each mutant strain, we assembled a phenotype profile composed of the fraction of cells with aberrant morphology for each of the 17 mutant classes, and computed the similarity of phenotype profiles between each pair of morphology mutant genes. Functionally related gene pairs exhibited significantly higher phenotype profile similarities, indicating that phenotype profiles were predictive of a functional relationship (Supplementary Figure 4a). Hierarchical clustering of phenotype profiles identified clusters enriched in functionally related genes, including clusters of genes involved in ER to Golgi transport, vacuole organization and exocytosis (Figure 4b). Interestingly, one cluster contained genes encoding regulators of actin and RNA splicing. Unlike most yeast genes, many actin regulatory genes, such as *COF1*, and *ARP2*, contain introns and thus depend on mRNA splicing to produce functional proteins and normal regulation of actin cytoskeleton organization (Figure 4b). The same cluster also includes the newly named *IPF1* gene (see above), additionally linking its function to actin cytoskeleton regulation.

Two poorly characterized genes, *YEL043W* and *NNF2* (*YGR089W*), had highly correlated phenotype profiles (PCC = 0.89) that were most similar to profiles of genes involved in Golgi vesicle and endosomal transport (Figure 4c, Supplementary Figure 4b). Both gene products are localized to the ER^20, 23, 24^ and contain coiled-coil domains that are often associated with vesicle tethering proteins^25^. Moreover, the coiled-coil domains of Yel043w and Nnf2 physically interact with each other^26, 27^ and the GI profiles of *YEL043W* and *NNF2* are both enriched for interactions with genes involved in vesicle trafficking^13^, suggesting these two proteins work together to promote vesicle trafficking. We named the *YEL043W* open reading frame *GTA1*, for **G**olgi vesicle **t**rafficking **a**ssociated.

In addition, many protein complexes that affect at least one of the screened endocytic markers had a high within-complex phenotype profile correlation (Supplementary Figure 4c, Supplementary Table 7). Phenotype profiles were more similar between components of the same protein complex structure that are in direct contact when compared to those that are not (Figure 4d). In some cases, these profiles were able to differentiate between closely related complexes, and between functional subunits of a complex. For example, ESCRT complex mutants led to the vacuolar class E and related phenotypes. Phenotype profiles were able to differentiate between ESCRT-I and ESCRT-II/III components (and to a lesser extent also between ESCRT-II and ESCRT-III components) (Figure 4e). In another example, phenotypic profiles differentiated two distinct functional subunits of the SPOTS complex, involved in sphingolipid homeostasis (Figure 4f). This modularity is consistent with the known biochemistry: the catalytic activity depends on Lcb1 and Lcb2, and is stimulated by Tsc3, whereas Sac1, Orm1, and Orm2 are believed to play regulatory roles (Figure 4f)^28^.

### Penetrance is an informative indicator of gene function

Besides specific phenotype information, an important output of our single-cell analysis was quantification of penetrance, defined as the total percentage of the population with an aberrant phenotype, in each mutant for each compartment. Among the morphology mutants were 1216 penetrance mutants that had a significant increase in penetrance compared to the control strain (Supplementary Table 2). For ∼90% of these mutants, the morphology defect was incompletely penetrant (Figure 5a). We binned mutants based on low, intermediate, or high penetrance and found that each group of genes was enriched for distinct functions (Supplementary Table 8). We previously showed that a network based on genetic interaction profiles provides a global view of the functional organization of the cell^13^. Thus, we next examined where these genes localized relative to biological process-enriched clusters on the global genetic interaction profile similarity network using spatial analysis of functional enrichment^29^ (SAFE) (Figure 5b). Highly penetrant mutants localized in close proximity to bioprocesses that are closely related to the function of the screened marker, genes corresponding to intermediate penetrance mutants mapped to “neighbouring” processes, and low penetrance mutants localized to clusters enriched for more functionally “distant” processes. For example, genes with highly penetrant Snf7-GFP phenotypes reflecting defects in late endosome morphology, mapped to clusters on the global genetic network representing multi-vesicular body sorting and vesicle trafficking, while genes exhibiting intermediate penetrance were located within vesicle trafficking-, glycosylation-, and polarity-enriched network clusters. Finally, low penetrance mutants tend to localize to regions of the global genetic network corresponding to vesicle trafficking, polarity, mRNA processing and transcription (Figure 5b).

**Figure 5:**
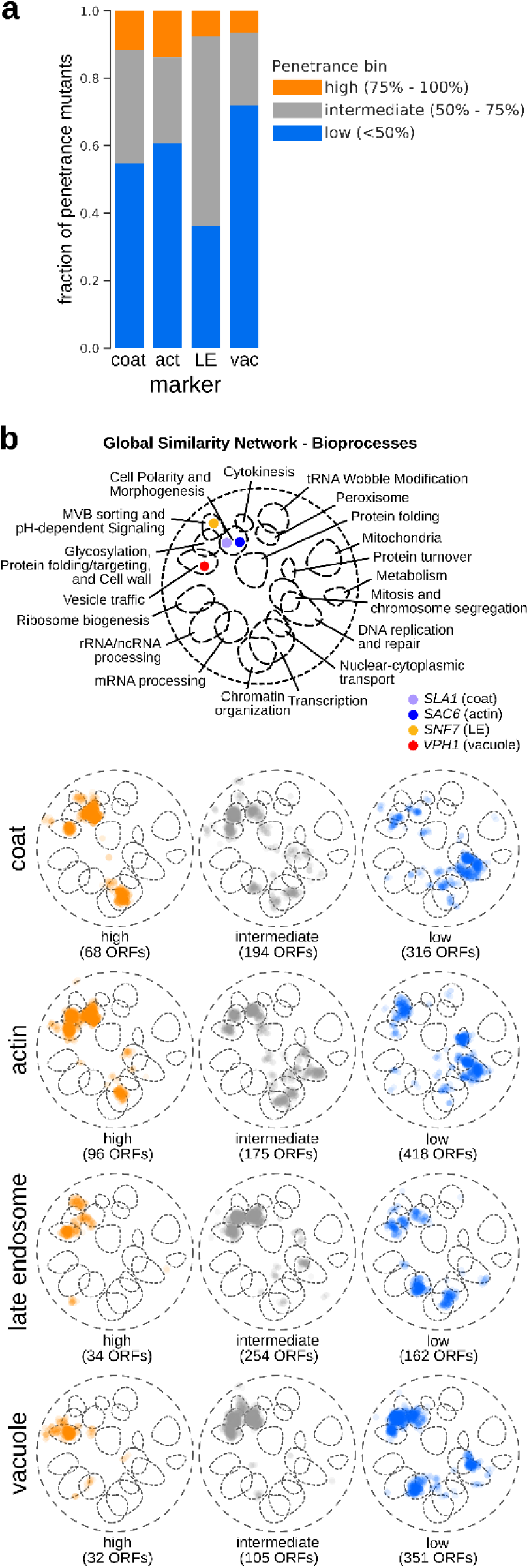
Functional analysis of incomplete penetrance. (see also Supplementary Table 8). (a) Stacked bar graph with fractions of penetrance mutants belonging to each penetrance bin for the four endocytic markers. act: actin; LE: late endosome; vac: vacuole. (b) SAFE (Spatial Analysis of Functional Enrichment) of penetrance mutants grouped according to penetrance. Top: Bioprocess key for interpreting the global similarity network for yeast genetic interactions visualized using SAFE, which identifies regions of the network enriched for specific biological processes (Costanzo et al., 2016). Coloured dots denote the localization of the 4 marker genes within the global similarity network. Below: SAFE of penetrance mutants grouped according to their penetrance and marker. Orange: genes whose mutation caused high penetrance; grey: intermediate penetrance genes; blue: low penetrance genes. Numbers in brackets refer to the number of unique ORFs in each group.

### Replicative age, asymmetric inheritance, and stress all contribute to incomplete penetrance in an isogenic cell population

Several factors have been suggested to affect penetrance in isogenic populations, including cell cycle position, cell size, replicative age, asymmetric segregation of molecular components, daughter-specific expression, and environmental factors^30–36^. Our quantitative single-cell analysis of the morphological defects associated with each marker provided a unique opportunity to explore the potential molecular and cellular mechanisms underlying penetrance.

#### Replicative Age and Penetrance

In yeast, replicative age can be assessed by staining chitin-rich bud scars to distinguish mother cells of different ages^37^. The average replicative lifespan for wild-type yeast (S288c) is 20-30 generations^38, 39^, thus old mothers are rare in a cell population. We examined wild-type control cells and five mutants with vacuole defects that had incomplete penetrance, including three mutants (*rrd2*Δ, *cka2*Δ, and *rpl20b*Δ) that are known to display a modestly extended replicative lifespan^39^, and two vacuole inheritance mutants^40^ (*vac8Δ* and *vac17Δ*). We stained and sorted the cells into bins roughly corresponding to number of bud scars, thus of unequal size, and assessed whether each cell had a vacuole defect.

For wild-type and all five mutants the fraction of outliers was lowest in the cells with lowest bud scar staining, corresponding to new daughters, and increased in mother cells with each cell division (Figure 6a, upper panel). In the bin with highest bud scar staining, corresponding to 5+ generations and consisting of ∼3% of the population, approximately half of the wild-type cells (53%) and from 51% to 94% of the mutant cells had a vacuolar morphology defect (Figure 6a-b). Thus, aberrant vacuolar morphology increases with the number of cell divisions even in young cells.

**Figure 6:**
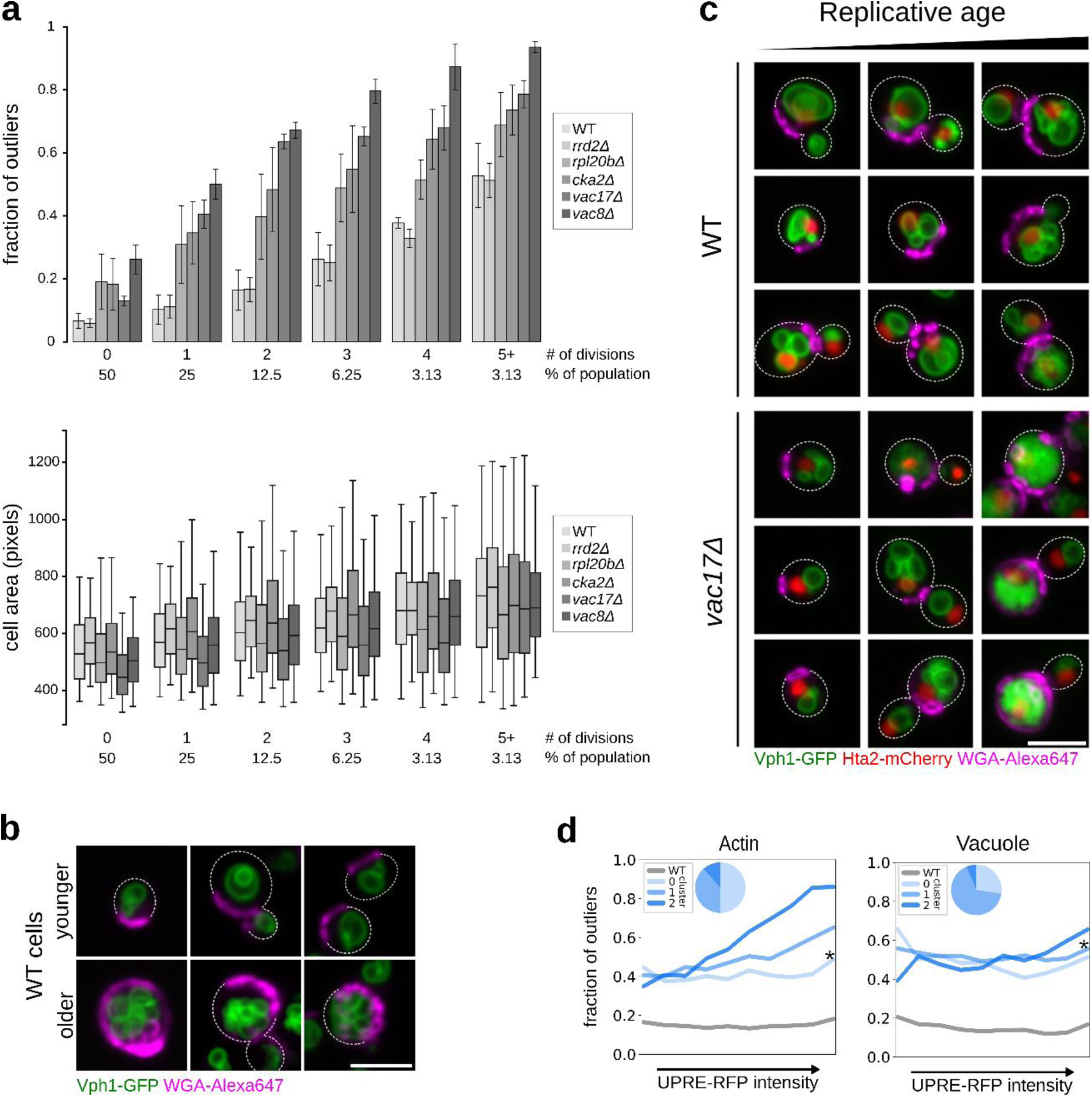
Factors contributing to incomplete penetrance. (see also Supplementary Table 8). (a) Penetrance as a function of replicative age. Top: Bar graph showing the fraction of outliers in populations of increasing replicative age (# of divisions) for wild-type (WT), and 5 mutant strains (*rrd2*Δ, *rpl20b*Δ, *cka2*Δ, *vac8*Δ and *vac17*Δ). Data are presented as mean of 3 biological replicates +/-SD. Bottom: Box plot with the distribution of cell sizes for the same populations of cells. Whiskers extend to the 5^th^ and 95^th^ percentile. (b) Micrographs of young (top row of images) and older (bottom row of images) wild-type (WT) cells expressing Vph1-EGFP (green vacuole) and stained with CF640R WGA (magenta bud scars). Scale bar: 5 µm. (c) Combined effect of replicative age and a vacuole inheritance defect on penetrance. Micrographs of wild-type and *vac17*Δ cells expressing Vph1-EGFP (green vacuole) and Hta2-mCherry (red nucleus), stained with CF640R WGA (magenta bud scars). Cells with increasing bud-scar staining (replicative age) are shown from left to right. Scale bar: 5 µm. (d) Relationship between stress response and penetrance. Single-cell UPRE-RFP levels were measured in ∼60 different mutant strains that we had identified as penetrance mutants with intermediate penetrance with defects in actin or vacuole morphology. Cells were binned into equal-sized bins, from low to high stress response, assessed as outlier or inlier, and clustered based on their penetrance profile (composed from the fraction of outliers in each stress-response bin). Each line plot represents a penetrance profile. * denotes the cluster with a profile most similar to wild-type. Insert pie charts show the proportion of mutant strains in each cluster.

Much of the work on replicative aging has been done on old mother cells but more recent studies have identified a number of factors that accumulate in relatively young mothers including oxidized proteins, protein aggregates and reactive oxygen species^34^. Multiple studies have reported that cell size increases in old mother cells^41, 42^. We quantified the size of our bud-scar-stained cells and confirmed that mother cells increased in size with replicative age, even in their first five generations (Figure 6a, lower panel), with no significant difference in cell size between wild-type cells and the mutants we assayed. Thus, in these experiments, increased penetrance seems to correlate with increased replicative age.

#### Asymmetric Organelle Inheritance and Penetrance

Organelle inheritance is an intrinsic component of cell division and mutations that affect this process can lead to cellular heterogeneity. In yeast, *VAC8* and *VAC17* are required for vacuole movement and partitioning between the mother and daughter cell^40^. We imaged cells of wild type and *vac17*Δ strains, with markers for vacuole and nucleus, stained for bud scars, and compared vacuole morphology defects in old and young cells of the two strains (Figure 6c). In these inheritance mutants, multilobed vacuoles were associated with aging and appeared at a much younger age compared to the wild-type background, leading to an increase in the fraction of the population that had a vacuolar morphology defect (Figure 6a, 6c). Thus, the observed cell-to-cell variability in deletion mutants of these two genes is a result of at least two factors: (1) defects in vacuole inheritance where daughter cells do not inherit a vacuole from their mother, but rather have to make one *de novo* (mother-daughter heterogeneity); and (2) replicative aging contributing to the accumulation of vacuole fission products with each cell division cycle, leading to multilobed vacuoles of increasing severity (replicative age-dependent heterogeneity). Similar to these vacuole mutants, asymmetric inheritance of many cellular components could affect penetrance.

#### Stress Response and Penetrance

Exposure to stress can lead to heterogeneous survival rates of isogenic yeast cells^35^, and can reduce penetrance in *Caenorhabditis elegans*^43^. Single cell analysis allowed us to address whether there was any relationship between levels of stress response and penetrance of morphology defects. We examined the unfolded protein response (UPR), which monitors folding of membrane and secreted proteins in the endoplasmic reticulum^44^. We first compared penetrance mutants with a study that had assayed UPR in the gene deletion collection using flow cytometry^45^. For actin and coat, an increased UPR was associated with mutants that had high penetrance in our screens (Supplementary Table 8). To explore the relationship between penetrance and the stress response in single cells, we crossed a reporter gene under the control of unfolded protein response elements^45^ (UPREs) into mutants that had incomplete penetrance for actin or vacuole defects (Supplementary Table 8). We then measured reporter activity as a proxy for the stress response level in each cell, divided by the cell area to normalize for cell size, and quantified penetrance as a function of stress response.

The relationships between penetrance and the UPR were different for the two assayed compartments, but the results were consistent with our correlation analysis (Supplementary Table 8). For approximately half of the mutants affecting actin, an increased UPR was associated with increased penetrance (Figure 6d, left panel, clusters 1 and 2), while the penetrance of vacuolar morphology was fairly constant across different levels of UPR for most mutants (Figure 6d, right panel, cluster 1). These findings indicate that UPR activation is correlated with penetrance of actin-based endocytosis phenotypes. At the molecular level, the UPR has been proposed to indirectly affect actin cytoskeleton remodelling by activating the cell wall integrity pathway^46–48^, which suggests that the connection between the UPR and actin-based endocytosis phenotypes may be causal.

## DISCUSSION

Here we describe a high-content screening pipeline that allowed us to interrogate sets of yeast mutants for effects on the morphology of the major endocytic compartments. Using a single-cell-level neural network classifier, we assigned over 16 million cells to one of 21 distinct endocytic phenotypes and obtained penetrance information for four markers for ∼5600 different yeast mutants (corresponding to ∼5300 genes, or ∼90% of the genes in the yeast genome). We found that ∼1600 unique yeast genes affect the morphology of one or more endocytic compartments. This dataset provides rich quantitative phenotypic information revealing roles of specific genes in shaping endocytic compartment morphology and the functional connections between genes and the compartments they perturb.

We used machine learning to perform outlier detection followed by classification of phenotypes to describe endocytic compartment morphology. These data allowed us to define possible morphologies for several functionally important cell compartments and also to build phenotype profiles, which summarize all assayed phenotypes associated with a specific genetic perturbation. The resulting phenotypic profiles predicted gene function and revealed functional information at the level of bioprocesses and protein complexes that was not evident by considering individual phenotypes.

Our analysis focused on markers that report on endocytosis, but the combined experimental and computational pipeline that we describe can be readily extended to other unrelated markers and phenotypes, enabling broader functional resolution. At this stage, the budding yeast system remains ideally suited to a large morphological survey of subcellular compartment morphology, given the availability of arrayed reagents for assessing loss- and gain-of-function perturbations in both essential and non-essential genes, and the ease of live cell imaging of strains carrying fluorescent markers^49, 50^. No matter the system used, a systematic analysis of phenotype profiles will greatly enhance our understanding of cellular function and lead to a more refined hierarchical model of the cell.

The rich phenotype information associated with single cell images enables the precise quantification of the prevalence of morphological phenotypes in a given cell population. We discovered that both incomplete penetrance, in which only a fraction of cells in a population have a mutant phenotype, and morphological pleiotropy, in which a specific mutation causes several phenotypically distinct subpopulations, are prevalent among mutant strains with defects in endocytic compartment morphology. More than half the morphology mutants we identified showed aberrant phenotypes for more than one of the four screened compartments, with the most pleiotropic mutants (those causing six or more specific phenotypes) being the most penetrant. Systematic analysis allows us to begin to explore the biological relevance and mechanisms of variable penetrance. For example, we were able to associate specific bioprocesses with high and low penetrance mutants, and to identify a number of protein complexes whose mutation is associated with morphological pleiotropy.

Many studies in mammalian cell systems have begun to address cellular heterogeneity using single-cell transcriptomics to identify sub-populations of cells in specific states, such as cancer, or during the cell cycle, cell differentiation, and exposure to stress^51–55^. Others have used cell imaging techniques to quantify both the structural and spatio-temporal properties of complex biological systems at the single cell level^56–59^. Regardless of the read-out, phenotypic heterogeneity appears to be a general feature of cell populations and so far, most studies have not directly addressed the biology underlying incomplete penetrance. Our ability to systematically assess single cell phenotypes in mutant cell arrays enabled us to show that replicative age, asymmetric organelle inheritance, and stress response all contribute to the incomplete penetrance of single gene mutations.

A number of other deterministic and regulated factors, such as noise in biological systems, micro-environment, epigenetic regulation, and the lipid and metabolic state of the cell have the potential to affect the penetrance and expressivity of a trait. In fact, for the majority of mutants, variability in morphological phenotypes between individual cells in an isogenic cell population is likely not driven solely by a genotype-to-phenotype relationship, but rather by a combination of smaller contributions from various effects that impact single cells differently depending on their physiological state. A deeper understanding of this variability may also have broad medical implications and should provide insight into the variable penetrance of genes affecting developmental programs^60, 61^ and disease genes^62, 63^.

## METHODS

### Query strain construction and construction of mutant arrays for imaging

To visualize endocytic compartments in living yeast cells, we C-terminally tagged 4 yeast proteins selected to visualize the endocytic compartments of interest with the yeast enhanced green fluorescent protein (yEGFP) or tdTomato. We used the polymerase chain reaction (PCR) to amplify an integration fragment containing: i) homologous regions 45 bp up- and down-stream of the target ORF’s C-terminus; ii) the fluorescent protein (FP) ORF and; iii) the selection marker. Plasmids pKT209^64^ (pFA6a-link-yEGFP-*CaURA3*), and pFA6a-link-tdTomato-*CaURA3* were used as templates. Plasmid pFA6a-link-tdTomato-*CaURA3* was constructed by replacing the yEGFP-*ADH1term* fragment between sites SalI/BglII in pKT209 with the tdTomato-*ADH1term* fragment. Switcher plasmid p4339 was used to exchange the *CaURA3MX4* cassette with the *NATMX4* resistance cassette to generate yEGFP-*NATMX4-* tagged strains^65^. Primers (starting with MMU_*) used to PCR FP-tagging cassettes for genomic integration are listed in the Supplementary Table 9. The lithium acetate transformation method^66^ was used to introduce the PCR product into yeast cells. The yeast proteins used as markers were: Sac6 for the actin module of actin cortical patches; Sla1 for the coat module of actin cortical patches; Snf7 for late endosomes; and Vph1 for vacuoles. All four proteins have been used previously as markers for these compartments^67–69^. *S. cerevisiae* strains and oligonucleotides used in the study are listed in the Supplementary Table 9.

To test for possible growth or other functional defects associated with the fluorescent protein tags, we performed the following tests: a) staining with FM 4-64 to check for a potential defect in endocytic internalization; b) real-time fluorescence microscopy imaging to check for potential fluorescent tag-effects on Sla1 and Sac6 endocytic patch formation dynamics; c) assessment of growth using serial spot dilutions on standard rich YPD media (1% (w/v) yeast extract, 2% (w/v) peptone, 2% (w/v) dextrose, 2% (w/v) agar) at different temperatures; d) mating of constructed FP-tagged query strains with strains carrying mutations in genes that had genetic interactions with *SAC6*, *SLA1*, *SNF7* or *VPH1*, followed by diploid selection, sporulation, and tetrad dissection to assess the growth of the double mutant progeny. A list of genetic interactions was obtained from^13, 70^. All of these experiments revealed no effect of the fluorescent tag on the tagged protein’s function, except for Snf7-GFP (and Snf7-tdTomato), where we confirmed an effect of the C-terminal fluorescent tag on Snf7p’s function, as has been observed previously with all ESCRT-III complex components^68^.

The constructed FP-tagged query strains were crossed to the haploid *MAT***a** deletion collection^12^ and to a collection of mutant strains carrying temperature sensitive (TS) alleles of essential genes^13, 14^. Haploid strains carrying both the fluorescent protein marker and the gene mutation from the mutant strain collections were selected using the SGA method^15^. All SGA selection steps involving a TS allele were conducted at permissive temperature (26°C). All SGA selection steps involving nonessential gene deletion mutants were conducted at 30°C. Sporulation was conducted at 22°C. For secondary, medium-scale screens, used also to determine penetrance reproducibility, false positive (FPR) and false negative rates (FNR), 1910 strains (36% of the complete array) were chosen from strains with both significant and non-significant phenotype fractions and SGA was done in biological duplicate. Strains included in the secondary array are marked in Supplementary Table 2.

### Preparation and imaging of live yeast cells

#### High-throughput microscopy

Yeast cell cultures were prepared for microscopy and imaged as previously described^23, 71^, with some modifications. Briefly, haploid mutant *MAT***a** strains expressing tagged FPs derived from SGA were grown and imaged in low fluorescence synthetic minimal medium^64^ supplemented with antibiotics and 2% glucose. Nonessential gene deletion mutants were grown and imaged in logarithmic phase at 30°C, and TS mutants of essential genes were first grown to mid-logarithmic phase and imaged at 26°C, and then incubated for three hours at 37°C and imaged at 37°C. Cells were transferred to a Concanavalin A (ConA) coated 384-well PerkinElmer CellCarrier Ultra imaging plate and centrifuged for 45 seconds at 500 rpm before imaging. To aid in cell segmentation, Dextran Alexa Fluor 647 (Molecular Probes) was added to cells in low fluorescence medium to a final concentration of 10 µg/ml before imaging.

For genome wide screens, micrographs were obtained on the Opera (PerkinElmer) automated spinning disk confocal microscope. Three fields with Z-stacks of 5 optical sections with 0.8 µm spacing were collected per well, with each field of view containing 50 - 150 cells. Secondary screens were imaged on an Opera Phenix (PerkinElmer) automated microscope. All imaging was done with a 60x water-immersion objective. Acquisition settings included using a 405/488/561/640 nm primary dichroic mirror. yEGFP was excited using a 488 nm laser, and emission collected through a 520/35 nm filter. tdTomato was excited using a 561 nm laser, and emission collected through a 600/40 nm filter. Dextran Alexa Fluor 647 was excited using a 640 nm laser, and emission collected through a 690/50 nm filter.

#### Monitoring the formation and progression of vacuolar class G phenotype with time-lapse fluorescence microscopy

Strains *his3*Δ (DMA1) and *sec18-1* (TSA54) from the *MAT***a** deletion and TS collections were crossed to strain Y15251. Haploid FP-tagged mutant clones were selected using the SGA method. Imaging plates were prepared as described above. Imaging was done using the Opera Phenix (PerkinElmer) automated system. Z-stacks of 5 optical sections with 0.8 um spacing were first acquired at room temperature, the temperature was then shifted to 37°C, and images were acquired at 1 h time intervals for 24 hours. Maximum z-projections, adjustment of intensity levels to optimize phenotype visualization, and image sequences were made with ImageJ^72^.

#### Assessing endocytic vesicle formation dynamics with live-cell imaging

Strains deleted for *YDL176W* (DMA754) or *HIS3* (DMA1; wild-type control) expressing Sla1-GFP and Sac6-tdTomato were grown to mid-log phase, immobilized on ConA-coated coverslips, and sealed to standard glass slides with vacuum grease (Dow Corning). Imaging was done at room temperature using a spinning-disc confocal microscope (WaveFX, Quorum Technologies) connected to a DMI 6000B fluorescence microscope (Leica Microsystems) controlled by Volocity software (PerkinElmer), and equipped with an ImagEM charge-coupled device camera (Hamamatsu C9100-13, Hamamatsu Photonics) and 100x/NA1.4 Oil HCX PL APO objective. Images were acquired continuously at a rate of 1 frame/second and analysed using ImageJ^72^. One hundred patches from 10-20 cells from two independent replicates were analysed per strain. Statistical significance was assessed with the unpaired t-test.

### Follow-up experiments related to the assessment of incomplete penetrance

#### Penetrance as a function of replicative age or vacuole inheritance

Strains *his3*Δ (DMA1), *rrd2*Δ (DMA4876), *rpl20b*Δ (DMA4693), *cka2*Δ (DMA4484), *vac17*Δ (DMA520), and *vac8*Δ (DMA1262) from the haploid *MAT***a** deletion collection were crossed to strain BY5841. Haploid mutants expressing the three FPs (*VPH1-GFP HTA2-mCherry* and *RPL39pr-tdTomato*) were selected using the SGA method. Cells were grown to logarithmic phase in standard conditions, washed in PBS, and stained with 400 µl 0.5 µg/ml CF640R wheat germ agglutinin (WGA) conjugate (CF640R WGA; Biotium) in PBS, nutating for 20 minutes at room temperature in the dark. Cells were then washed 3x with PBS, placed in low fluorescence medium, and transferred to a ConA treated imaging plate. Acquisition of z-stacks was done on the Opera Phenix (PerkinElmer) automated microscope as described above. Maximum z-projections, channel merging and adjustment of intensity levels to optimize subcellular signal visualization (used only for figures) were made with ImageJ^72^. The experiment was done in biological triplicates.

#### Effect of the UPR pathway

A *URA3*::UPRE-mCherry cassette, which encodes mCherry driven by a minimal *CYC1* promoter and four tandem unfolded protein response elements (UPREs), was amplified using PCR from pPM47^73^ and integrated at the *URA3* locus in BY4741. Primers used (URA3pr-F and dn_mCherry-R) are listed in the Supplementary Table 9. Plasmid pPM47 was a gift from Feroz Papa. The strain with integrated UPRE-mCherry was crossed to query strains containing *SAC6-GFP::NATMX4* or *VPH1-GFP::NATMX4* and tetrads were dissected to obtain query strains with a GFP-tagged morphology marker and UPRE-mCherry (strains BY6279 and BY6285).

A mini-array of gene-deletion strains identified as intermediate penetrance mutants for actin was chosen and crossed to BY6285. Likewise, a mini-array of vacuole mutants was crossed to BY6279. SGA was used to select haploid strains with both the marked morphology compartment and the stress reporter. Cells were grown for imaging using standard conditions and imaged in low fluorescence medium containing 5 µg/ml Dextran Alexa Fluor 647 on an Opera Phenix (PerkinElmer) automated system as described above.

### Determining Single Mutant Fitness (SMF) of the DMA-SLOW collection

In order to determine the single mutant fitness for slow-growing nonessential gene deletion strains (DMA-SLOW collection), that were previously excluded from the global genetic interaction analysis^13^ (∼400 ORFs), we carried out 5 SGA screens where a WT query strain carrying a *NATMX* marker inserted at a neutral locus (Y8835) was crossed to the *KANMX*-marked DMA-SLOW collection. SGA screens were performed at 30°C. Colony size was quantified using SGATools^74^.

### Image analysis and object quality control

#### Image pre-processing, object segmentation and quantitative feature extraction

Acquired stacks were compressed into a maximal z-projection using ImageJ^72^. CellProfiler^16^ was used for object segmentation and quantitative feature extraction. Cells were segmented from intensity-inverted Dextran Alexa Fluor 647-channel images. Cell intensity measurements of the Dextran Alexa Fluor 647 channel were collected for quality control purposes. Segmented cell boundaries were then applied to the endocytic marker channel to segment secondary objects (endocytic compartments), define tertiary objects (cytoplasm), and extract area, shape, intensity, and texture measurements of the segmented endocytic compartments, cytoplasm and whole cell. Two additional features were calculated from the extracted CellProfiler features: a) fraction of the cell occupied by the screened compartment(s) (compartment_areashape_area divided by cell_areashape_area); and b) compartment diameter ratio (compartment_areashape_maxferetdiameter divided by compartment_areashape_minferetdiameter). In total, we extracted quantitative information for approximately 21 million single cells, and approximately 73 million individual endocytic compartments. The raw data were imported into a custom-made PostgreSQL database.

#### Cell quality control

To reduce noise in the analysis due to segmentation artifacts and ensure only high-quality objects were included in downstream analyses, a quality control filter was applied to all segmented cells. First, the quality control filter discarded low-quality cell objects based on shape, size and intensity measurements collected from the Dextran Alexa Fluor 647 signal. These low-quality objects included badly segmented cells, ghost objects (segmented background), clumps of cells, and dead cells. Second, all images with < 5 cells were excluded. Third, all wells with only 1 good site (out of the 3 acquired) were excluded. Additionally, we trained a 2-layer fully connected neural network to identify and exclude from the dataset small buds that had been segmented independently of mother cells, and bad cells missed by other quality control filters (see below for details). Across all screens, the used filters discarded 20% of cell objects, leaving approximately 16.3 million good cells for subsequent analysis. On average, 640 good cells for each strain from 2.6 biological replicates were retained for downstream analyses.

### Data processing, outlier detection, and classification

#### Data pre-processing

The extracted features were standardized by computing mean and standard deviation of features from wild-type control strains (negative controls) to remove feature means and scale to unit variance separately for each imaged 384-well plate in a screen. Means and standard deviations of features from each imaged plate were analysed to identify potential batch effects on plates.

#### Selection of positive controls

Positive control mutants were selected based on phenotypes annotated in the *Saccharomyces* genome database (SGD, https://www.yeastgenome.org) and published literature (Supplementary Table 1). Only mutants for which we were able to visually confirm the published phenotype in our images were included in the positive control set. Additionally, to ensure all main phenotypes were included in our classifier, an unsupervised outlier detection approach was used to search for mutants with unpublished or poorly annotated phenotypes (see below for details). The two approaches combined gave us a set of 21 different subcellular morphologies, comprising 4 wild-type and 17 mutant phenotypes. We note that our vacuole phenotypes do not perfectly overlap with the vacuolar morphological classes that have been described previously^75^ (classes A to F). To avoid confusion, we adopted descriptive names for most of our vacuolar phenotype classes (Figure 1c, Figure 2).

The lists of positive control strains associated with each mutant phenotype were subsequently used to compile the classifier training set (see below for details). Visual inspection of all micrographs from positive control mutants was used to assign each mutant to a penetrance bin (100 - 80%, 80 - 60%, 60 - 40%, 40 - 20%, 20 - 0%). These manual penetrance assignments were used to validate the accuracy of computational penetrance assignments obtained through classification (see below for details). Positive control mutants and their manual penetrance assignments are listed in Supplementary Table 1.

#### Unsupervised outlier detection

An unsupervised outlier detection approach was used to identify additional positive control strains (see *Selection of positive controls*). First, principal component analysis (PCA) was applied to the extracted CellProfiler features to reduce the redundancy and correlation of features in the data^76^. The number of PCs was selected so that at least 80% of the variance in the complete data was explained. Next, to identify mutant strains that affect the morphology of the imaged subcellular compartment, an outlier detection method was implemented with the goal of detecting cells whose morphology differed substantially from the negative (wild-type) controls. In the feature space, we identified cells with non-wild-type morphologies based on their distances from the negative control distribution. To quantify these distances, we implemented a one-class support vector machine (OC-SVM)-based outlier detection method^77^. We used OC-SVM implemented in Python’s scikit-learn package with default hyper-parameters (radial basis function kernel, kernel hyper-parameter gamma set to 1/N where N is the number of used PCs, and hyper-parameter nu set to 0.5 in order to define a stringent population of negative control cells). For each single cell in the complete dataset, we calculated the distances to OC-SVM decision function. Next, we applied a threshold on the calculated distance at the 20^th^ percentile of the negative control cells to differentiate in- and outliers.

For each mutant strain, unsupervised penetrance was defined as the percentage of outlier cells obtained from the unsupervised outlier detection approach. The statistical significance of penetrance was calculated using a hypergeometric test (identical to one-tailed Fisher’s exact test)^78^, with negative controls as the background. For each endocytic marker, top scoring mutant strains were visually inspected to identify any major mutant phenotypes that had been previously missed, and additional positive control strains (see above).

#### Single cell labelling and classification

##### Single cell labelling tool

The custom-made single cell image viewer is a Django-based web application written in Python which allows the user to input different parameters or filters and then view the cells satisfying the set conditions. The web interface is developed using HyperText Markup Language (HTML 5), Cascading Style Sheets (CSS3) and JavaScript. Taking advantage of Django’s capability to use multiple databases, the primary PostgreSQL database containing raw CellProfiler features and unique cell IDs was used in this tool to pull the information needed to display each single cell. The information needed was: the image to which the cell belongs, the image’s location on the server, and the x- and y-coordinates of the cell. The tool allows the user to label and save a phenotype for a specific cell which would then be saved to the single cell viewer database and used to compile the cells’ features for the training set.

##### Manual labelling of single cells

For each marker, and separately for the primary genome-wide and secondary medium-scale screens, single cells from positive control mutant strains as well as negative control wild-type strains were manually assigned to a mutant or wild-type phenotype class using the single cell labelling tool. In total, 42 sets of labelled cells were compiled (2 types of screens x 21 phenotypes; i.e. 4 wild type and 17 mutant phenotypes).

For cell quality control purposes (see *Cell quality control* above), for each Sla1 and Sac6 screen, we manually labelled approximately 320 small buds, that had been segmented independently of their mother cells (“small bud” class), and approximately 250 badly segmented cells (“bad” class). We trained a 2-layer fully connected neural network (see below for details) with these two cell quality control classes and the wild-type class to identify small buds that had been segmented independently of mother cells, and bad cells missed by other quality control filters. All cells that were assigned to the “small bud” or “bad” classes with an average prediction probability across 10 random initializations of >=85% were excluded from the final set of good cells.

##### Training set clean-up

To identify cells in the training set that were mislabelled, we did an initial training run with all of the labelled positive and negative control cells, as described above. For training, 5-fold cross–validation on the labelled dataset was used. Each fold was split into 80% for training set (20% of training set is used for validation set) and 20% for test set during neural network training. Each fold was used for training a fully connected 2 hidden-layer neural network (2NN) 10 times with different random initializations. With this approach on 5-fold cross-validation and 10 random initializations, we obtain 10 separate predictions for each of the labelled cells. Any cells that were incorrectly classified in two or more random initializations were manually inspected and cells that were originally mislabelled were removed from the final labelled set. Up to 12% of the labelled cells were removed from each training set using this approach. All subsequent training of the 2NNs was on this cleaned dataset. The final number of labelled cells in each phenotype ranged from 35 to 982, with an average of 420 cells.

##### Visualization of the training set feature space with t-SNE

To assess whether the extracted CellProfiler features could be used to accurately distinguish between different phenotypes, the high-dimensional feature space for each of the single cells from the training sets was visualized using a non-linear dimensionality reduction technique - t-distributed stochastic neighbour embedding^17^ (t-SNE). Python’s scikit-learn package (version 0.19.0) was used for t-SNE with default hyper-parameter settings except for perplexity. The perplexity hyper-parameters chosen for the 4 markers were: 50 for Sac6, 30 for Sla1, 60 for Snf7, and 50 for Vph1.

##### Classification: single-cell level assignment of mutant phenotypes

To classify all cells from the final dataset (see *Cell quality control* above) into different mutant phenotypes, the training sets comprised of labelled single cells were used to train a fully connected 2 hidden-layer neural network (2NN). We trained a 2NN for each of the endocytic markers and screen types, totalling eight trained 2NNs. We opted for separate training sets for the two screen types (genome-wide and secondary screen), as this strategy gave us better classification accuracy (possibly because the two screen types were imaged using different microscopes). The 2NN was implemented using Keras (https://keras.io) with TensorFlow backend (https://tensorflow.org).

The input layer consisted of the scaled single-cell CellProfiler features and we used a soft-max output layer^79^. The first hidden layer had 54 units and the second hidden layer had 18 units. The hidden layers used ReLU activation functions^80^. All the hyper-parameters used in the training for the Stochastic Gradient Descent optimizer^81^ are specified at https://github.com/BooneAndrewsLab/ODNN. We used the same architectures and hyper-parameter settings for each network; these hyper-parameters were selected to provide good performance without overfitting, with a short training period on the whole unfiltered training sets.

For training, 5-fold cross–validation on the labelled dataset was used. Each fold was split into 80% for training set (20% of training set is used for validation set) and 20% for test set during neural network training. Each fold was used for training a 2NN 10 times with different random initializations, resulting in 10 predictions for each cell in the test set. The final probability of each cell in the test sets was calculated by averaging the probabilities from these 10 randomly initialized 2NNs. The class with the maximum average probability was used for the predicted label. The combined test set predictions were displayed in a confusion matrix and used to assess the neural network’s performance. Most phenotypes could be classified with very high accuracy, except those with the smallest training sets (Supplementary Figure 1c).

The 2NN employs a relatively new strategy for creating an ensemble classifier; to ensure that this strategy did not create bias in its classifications, we compared it to two more traditional approaches to creating an ensemble classifier. Specifically, 10 base classifiers in the ensemble differed only in their random initializations but shared hyperparameters and training sets. This strategy permits us to use the entire training set to train each classifier rather than only the ∼68% used in bagging. For classifiers with multiple local optima, such as neural networks, the new strategy has shown better performance generalization and uncertainty calibration than bagging^82^. However, to validate this method in our context, we compared it to two other approaches using the Sac6 and Vph1 genome-wide screens. One approach (2NN – CVx1) employed the same training strategy as 2NN, but predictions of unlabelled cells in the full dataset were done during each of the 10 random initializations and 5-fold cross-validations. In other words, we averaged the output of 50 networks trained on five different, partially overlapping, training sets. The second approach (2NN – CVx10) did not include 10 random initializations, and the training was done using 10 independent runs of 5-fold cross-validations. Here, we averaged the output of 50 networks trained on 50 different, partially overlapping, training sets. Similar to 2NN – CVx1 approach, with this third approach, we predicted the entire dataset of unlabelled cells during each of the 10 independent runs and 5-fold cross-validations. We assigned cells in a screen the phenotype with the highest average probability across the 50 2NNs for both 2NN – CVx1 and 2NN – CVx10. The average correlation between these three approaches on penetrance values and phenotype fractions across all genes in the two genome-wide datasets was 0.95 (Supplementary Table 1). We thus concluded that our use of the same training sets and hyperparameters in the ensemble in our 2NN classifier did not introduce biases compared to a scheme which employed different training sets on each classifier.

After estimating the general performance of the 2NNs using cross-validation, to make predictions on the entire dataset of unlabelled cells, we retrained the networks on the entire filtered training set. Similar to the approach described above, we trained 10 separate 2NNs starting from different random initializations and assigned cells in a screen the phenotype with the highest average probability across the 10 2NNs. Mean single-cell prediction probabilities are included in Supplementary Table 1. The 2NN classifier assigned the highest classification probabilities to those cells that were most similar to those in the manually labelled training sets (Figure 2), but at the same time allowed us to correctly assign cells with different severity of a particular phenotype to the same class.

Additionally, to identify any strains with phenotypes not included in our classifiers, we assigned all cells with low classification probabilities to a “None” class. Cells were assigned to the “None” class when the maximum probability was lower than 2/N (where N = number of phenotypes). Visual inspection of strains with the highest fraction of cells assigned to the “None” class for each marker revealed no additional phenotypes (Supplementary Table 2). This approach does not exclude the possibility that cells with additional rare or non-penetrant phenotypes were incorrectly classified. For example, we did not include the class F phenotype (large central vacuole surrounded by several fragmented vacuolar structures) in our vacuole classifier, since none of the previously reported mutants had significant fractions of the population displaying the phenotype^75^. The class F cells were therefore classified as wild-type, enlarged or multilobed.

All cells assigned a non-WT phenotype were defined as outliers. For each strain, the fraction of the cell population displaying each phenotype (specific phenotype fraction), as well as the penetrance (defined as: penetrance = 1 - % WT cells) were calculated. The phenotype fractions and penetrance calculated from 2NN classification were used in all subsequent analyses.

Custom scripts for data pre-processing, running the supervised 2 hidden-layer fully connected neural network for single cell classification, and penetrance calculation are available at: https://github.com/BooneAndrewsLab/ODNN.

### Penetrance reproducibility

#### Estimating penetrance reproducibility

To assess the quality of our computed penetrance values, we first compared them to visually assigned penetrance estimates and found a strong agreement for all four screened markers (average Pearson correlation coefficient (PCC) = 0.87) (Supplementary Figure 1d, Supplementary Table 1). We next assessed penetrance reproducibility by determining: (i) the difference in the calculated penetrance; and (ii) the Pearson correlation coefficient (PCC) between replicates of mutant strains for each marker and screen type. Across all screens and markers, the average PCC between replicates is 0.65 (two-tailed p-value = 0). Finally, we focused on the replicate pairs with a penetrance difference > 30, and determined the prevalent causes leading to penetrance irreproducibility. Cell number had the biggest impact on penetrance reproducibility, since replicates with low cell counts represented 60% of replicate pairs with highest penetrance differences (Supplementary Figure 1e). Among low cell count replicates, 2/3 were from strains with considerable growth defects, making single mutant fitness (SMF) a good indicator of penetrance reproducibility. For the remaining 40% of replicates with a large penetrance difference, 42.6% could be attributed to temperature sensitive strains (35.2% of large difference replicates) and technical artifacts (cross-contamination, bad segmentation, failed quality control, or misclassification) (7.4% of large difference replicates), while for 57.4% of replicates we could not identify a clear cause. The increased penetrance difference between biological replicates of TS strains could be a consequence of small differences in growth and imaging temperature between replicates.

#### Bootstrapping to determine sufficient cell count

Since different strains and plates varied greatly in the number of imaged cells, we used a bootstrapping approach to determine the standard deviation between replicates of varying cell counts and to estimate the minimal cell number required to obtain a confident penetrance calculation (Supplementary Figure 1f). Increasing numbers of cells were sampled from every screen individually, screens combined by marker, and all screens combined. Sampling was done on two scales: first on a small scale ranging from 10 to 100 cells in increments of 10, then on a larger scale ranging from 125 cells to 1000 cells in increments of 25. Cell sampling was done one hundred times from populations of approximately 200,000-670,000 cells (for individual screens), and the average penetrance and standard deviation of the 100 independent samplings for each sample size were calculated and plotted (Supplementary Figure 1f). Based on the wide distribution of wild-type replicate penetrances (Supplementary Figure 1g), we chose a relative standard deviation of 0.2 (which is equal to approximately +/-4 penetrance points for a wild-type population) as our confidence threshold. The average minimum required cell number across different markers and screens was 98 cells, and this criterion was satisfied by 83.4% of imaged samples. We next examined the potential impact of cell-density effects, such as gradients, on penetrance, and observed no significant effects (Supplementary Figure 1h). R^2^ between cell number and calculated penetrance for all replicates that met the minimum cell count was 0.0047.

### Identification of morphology mutants, calculation of accuracy, false positive and false negative rates

#### Specific phenotype mutants (SPMs)

For each phenotype (17 mutant phenotypes representing 4 for patch actin module, 4 for patch coat module, 3 for late endosome, 6 for vacuole) and screening condition (room temperature, 30°C, or 37°C), the threshold (thr) for the specific phenotype fraction was defined as the phenotype fraction value corresponding to the 98^th^ percentile of the distribution of the specific phenotype fraction across all wild-type replicates. Since each 384-well imaging plate had wild-type strains at 76 positions (all border wells), a full genome-wide screen had more than 1800 wild-type replicates. In cases where this calculated threshold was < 0.05, the threshold was set to 0.05. Additionally, the stringent threshold for the fraction of a specific phenotype was defined as: str = (max - thr) x 0.25 + thr; where max is the highest observed penetrance for that phenotype.

The final specific phenotype fraction for each mutant strain was calculated from the genome-wide and secondary screen values as the replicate number-weighted mean phenotype fraction. Each mutant strain had to satisfy the following criteria in order to be considered an SPM: a) weighted mean phenotype fraction >= phenotype fraction threshold (or stringent threshold for stringent SPMs); b) >=50 good cells and; c) >=10 cells assigned with the phenotype in question. SPMs, stringent SPMs and thresholds are listed in Supplementary Table 2.

#### Penetrance mutants

For each marker (Sac6, Sla1, Snf7, and Vph1), and screening condition (room temperature, 30°C, or 37°C) the penetrance threshold was defined as the penetrance value corresponding to the 95^th^ percentile of the penetrance distribution of all wild-type replicates. The final penetrance for each mutant strain was calculated from the genome-wide and secondary screen values as the replicate number-weighted mean penetrance. Each mutant strain had to satisfy the following criteria in order to be considered a penetrance mutant for a given marker: a) weighted mean penetrance >= calculated penetrance threshold; b) >= 50 good cells; c) Bonferroni corrected p-value < 0.05 (for strains not included in the secondary array). Penetrance mutants and thresholds are listed in Supplementary Table 2.

The specific phenotype and penetrance mutant groups comprise 1623 yeast genes, of which 66.5% (1079) were both SPMs and penetrance mutants, 25.1% were only SPMs (407), and 8.4% (137) were only penetrance mutants (also referred to as non-phenotype-specific mutants; see Figure 1d). In general, the ORFs that qualified as only non-phenotype-specific mutants or only SPMs were either (i) SPMs with significant but smaller mutant phenotype fractions that did not qualify as penetrance mutants, or (ii) non-phenotype-specific mutants with one or more mutant phenotypes with fractions below the specific phenotype significance thresholds. For example, a deletion mutant of *RRD2*, which encodes a component of a serine/threonine protein phosphatase involved in Tor1/2 signalling, had 42% of cells with aberrant vacuolar morphology, which is above the ∼32% penetrance threshold for the vacuolar marker. However, none of the six specific vacuolar phenotype fractions exceeded the respective SPM thresholds (see Supplementary Table 2 for details).

We note that although a number of TS strains displayed increased levels of non-wild-type-looking cells even at room temperatures (data available at thecellvision.org/endocytosis), consistent with previous work^14^, we used only data from TS strains grown at 37°C for the identification of morphology mutants and downstream analyses.

#### Accuracy, FPR, FNR

The accuracy, false positive (FPR) and false negative (FNR) rates were calculated from biological replicates (same query strain, same screening condition, same microscopy setup) of mutant strains as follows:

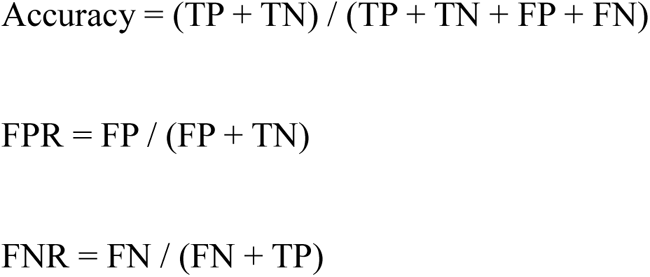

A replicate pair was called a true positive (TP), if both measurements satisfied the criteria for penetrance mutants described above (penetrance > threshold, >= 50 cells, and corrected p-value < 0.05). Similarly, a replicate pair was called a true negative (TN) when neither of the two replicates satisfied the criteria for penetrance mutants. False positives (FP) and false negatives (FN) were those pairs where one replicate was a penetrance mutant, while the other was not. The estimated average accuracy was 86.6%. The estimated false positive rate was 9.5%. We estimate that the false positives are mainly found in the intermediate penetrance range. The false negative rate was higher at ∼24.9%, as expected from a stringent cut-off.

#### List of consensus morphology mutants

A consensus rule for genes with multiple screened alleles was used. For each marker and phenotype, a gene was considered a penetrance mutant or SPM if half or more of its alleles satisfied the respective significance criteria. Consensus morphology mutants are listed in Supplementary Table 2 (labelled as consensus_*).

For penetrance bin-dependent analysis, for genes with multiple alleles and with a penetrance mutant in the consensus list, we defined the penetrance as the maximum penetrance among the screened alleles that qualified as a penetrance mutant (see above). Penetrance bins of all consensus morphology mutants are listed in Supplementary Table 2.

### Enrichment and correlation analyses

#### Data standards used in the analyses

##### Protein complex standard

The protein complex standard was downloaded from the EMBL-EBI Complex Portal (https://www.ebi.ac.uk/complexportal/home) and is included in Supplementary Table 4.

##### Gene ontology biological process standard

Biological process categories for functional enrichment were derived from a standard set of GO Slim biological process term sets downloaded from the Saccharomyces Genome Database (www.yeastgenome.org/).

##### Biological pathway standard

The pathway standard was downloaded from the KEGG database^83^ and is included in Supplementary Table 4.

#### Gene features used in the analyses: Numeric features

##### Marker abundance data

For each mutant strain, the mean FP intensity (CellProfiler feature cell_intensity_meanintensity) extracted from the genome-wide screens was used to calculate the relative marker abundance (“Marker relative abundance”) and relative standard deviation of marker abundance (“Marker abundance CV”). For each mutant strain, the calculations were normalized to the per-plate wild-type strain values. The relative marker abundance and marker abundance CV data are included in Supplementary Table 4.

##### Cell size data

For each mutant strain, raw single-cell size data (corresponding to cell area in pixels; CellProfiler feature cell_areashape_area) extracted from the genome-wide screens was used to calculate the relative mean cell size (“Relative cell size”) and relative standard deviation of cell size (“Cell size CV”). For each mutant strain, the calculations were normalized to the per-plate wild-type strain values. The relative cell size and cell size CV data are included in Supplementary Table 4.

##### Single mutant fitness

Single mutant fitness (SMF) values for nonessential gene deletion strains (DMA), and essential gene temperature-sensitive strains (TSA) were taken from Costanzo et al., 2016^13^. For the slow-growing nonessential gene deletion collection (DMA-SLOW), mean colony size measurements from 5 wild-type SGA screens were used to estimate single mutant fitness. Colony size was quantified using SGATools^74^. SMF was calculated as the relative colony size compared to wild-type^84^. The SMFs of all strains used in this study are listed in Supplementary Table 2.

##### Broad conservation

Broad conservation is a count of how many species, out of a set of 86 non-yeast species, have an ortholog of a given gene. Broad conservation was assessed as described in Costanzo et al., 2016^13^.

##### Positive, negative, and total number of genetic interactions

The numbers of positive, negative and all genetic interactions (“Positive GI/Negative GI/Total GI”) for each mutant strain were extracted from TheCellMap^85^ (www.thecellmap.org). For genes with multiple alleles, the number of GIs was averaged across alleles.

##### PPI degree

Protein-protein interaction data was retrieved from BioGRID^86^ and refers to the union of five high-throughput studies^87–91^.

##### Pleiotropy

Pleiotropy data were from Dudley et al., 2005^92^. The number of conditions (out of 22 tested) that lead to reduced fitness was used as a measure of pleiotropy.

##### Multifunctionality

The total number of annotations across a set of functionally distinct GO terms described in Myers et al., 2006^93^ was used as a multi-functionality index. Multifunctionality was assessed as described in Costanzo et al., 2016^13^.

##### Phenotypic capacitance

The phenotypic capacitance was used directly from Levy and Siegal^94^ and captures variability across a range of morphological phenotypes upon deletion of each of the nonessential genes.

##### Co-expression degree

This measure is derived from a co-expression network based on integration of a large collection of expression datasets^95^. Co-expression degree was assessed as described in Costanzo et al., 2016^13^. Pairs of genes with a MEFIT value above 1.0 were defined as co-expressed.

##### Expression level and transcript counts

The expression level values reflect the mRNA transcript levels of all yeast genes in wild-type cells grown in YPD measured using DNA microarrays^96^. Transcript counts indicate the number of mRNA copies of each transcript per cell^97^.

##### Molecules per cell

Protein abundance data was derived from the unified protein abundance dataset compiled from 21 quantitative analyses^98^. The “mean molecules per cell” values were used for analysis.

##### Expression variance measured under different environmental conditions

For each gene, the variance in expression across all conditions surveyed in Gasch et al., 2000^99^, was measured. This dataset contains yeast gene expression levels measured in response to a number of different environmental conditions. For details refer to Costanzo et al., 2016^13^.

##### Protein abundance and localization variation

Data on protein abundance variation (“Protein abundance RV”), and subcellular spread (“Subcellular spread RV”) were from Handfield et al., 2015^100^.

##### UPRE level

Data for UPR-levels was from Jonikas et al., 2009^45^.

#### Gene features used in the analyses: Binary features

##### Whole genome duplicates

This binary feature reflects whether each gene has a paralog that resulted from the whole genome duplication event^101^.

#### Other datasets

##### Endocytic internalization dataset

Data on endocytic internalization levels in nonessential gene deletion mutants was from Burston et al., 2009^8^. Deletion strains with an invertase score below the median (no assigned value in the published dataset) were assigned a value of 0.

##### Orthologs

A set of *Homo sapiens* orthologs of *S. cerevisiae* were obtained from the InParanoid eukaryotic ortholog database version 8.0 (http://inparanoid.sbc.su.se).

##### Essential and Nonessential Gene Sets

The essential and nonessential gene lists were obtained from Saccharomyces Genome Database (SGD; www.yeastgenome.org/).

#### Morphology mutant enrichment analyses

##### GO Slim biological process

We performed the GO biological process enrichment analysis for each set of SPMs using the GO Slim mapping file available through the Saccharomyces Genome Database (www.yeastgenome.org/). SciPy’s hypergeometric discrete distribution package was used to calculate p-values. P-values were adjusted using the Bonferroni correction. Fold enrichment was calculated as (mutants in term / all mutants) / (term size / all background).

##### Protein complex and biological pathway

For each phenotype, we calculated the number of mutants that coded for members of each protein complex and tested for enrichment by using one-sided Fisher’s exact tests. To identify specific enrichments associated to phenotypes, and not associations caused by genes that were morphology mutants in many phenotypes, we randomized the phenotype-gene associations. Then, for each randomized network, we calculated the number of morphology mutants that belonged to each complex. We only reported phenotype/complex enrichments with a Fisher’s p-value (see “p_greater”) below 0.05 and with phenotype/complex overlaps in the real network (see “P1”) higher than 95% of the overlaps observed in the randomized networks (see “p_rnd”). The same approach was used to evaluate morphology mutant enrichment of KEGG biological pathways. Used standards are included in Supplementary Table 4. Enrichment results are included in Supplementary Table 3.

##### Gene feature

We compared the values of morphology mutants and genes not identified as morphology mutants against a panel of gene features. We computed statistics by performing one-sided Mann-Whitney U tests for numeric features (p-value < 0.05), and by one-sided Fisher’s exact tests for binary features (p-value < 0.05). For features with data for multiple alleles, values for different alleles were averaged. For each numeric feature, we performed a z-score normalization in which we used the median (instead of the mean) and standard deviation of non-morphology-mutants. Since all median z-scores of the non-morphology-mutant sets were centred to zero, in plots we reported only the median z-score values for morphology mutants. For each binary feature, we calculated the fraction of morphology mutants (f_hits) and non-morphology-mutants (f_nonhits) with that particular feature. Then, we calculated the fold enrichment as the logarithm of f_hits divided by f_nonhits.

We followed the same approach to compare: i) morphology mutants and non-morphology-mutants of each individual marker (vacuole, late endosome, coat, and actin); ii) penetrance mutants with high (>= 75%), intermediate (75% > x >= 50%), and low (< 50%) penetrance values vs non-morphology-mutants for each individual marker; iv) SPMs vs genes that were not morphology mutants for each of the 17 mutant phenotypes. Results of these analyses are provided in Supplementary Table 3 and Supplementary Table 8.

46 ORFs that are present in the screened array, but have been deleted from SGD, were excluded from analysis (list of excluded ORFs is available at thecellvision.org/endocytosis).

#### Comparison of direct and indirect protein contacts

We used Interactome3D^102^ (version 2019_01) to select available protein complex structures in the PDB with three or more yeast proteins, and identified which of the proteins in the complex were in direct contact. Interactome3D defines direct contacts between two proteins if they have at least five residue-residue contacts, which can include disulphide bridges (i.e., two sulphur atoms of a pair of cysteines at a distance ≤2.56 Å), hydrogen bonds (i.e., atom pairs N-O and O-N at a distance ≤3.5 Å), salt bridges (i.e., atom pairs N-O and O-N at a distance ≤5.5 Å), and van der Waals interactions (i.e., pairs of carbon atoms at a distance ≤5.0 Å). We classified proteins in the same complex structure that did not meet our criteria for direct contact as indirect contacts. Additionally, we compiled a list of protein pairs belonging to different protein complex structures.

For each screened gene, we built a profile with its 17 specific phenotype fractions (phenotype profile; phenotype fraction data is provided in Supplementary Table 2). For genes with several screened alleles, we used the mean specific phenotype fraction across alleles. Strains with incomplete profiles (missing data for any of the 4 markers) were excluded from the analysis. For each pair of profiles, we calculated their Pearson’s correlation.

Correlation values were then grouped by the relationship of proteins in experimental structures: i) protein pairs in contact in a protein complex structure; ii) protein pairs in the same experimentally solved protein complex structure but not in direct contact; or iii) protein pairs that do not belong to the same solved protein complex structure. Difference between the sets of correlation values was evaluated by one-sided Mann-Whitney U tests. Files with gene-averaged specific phenotype profiles, and profile PCCs are available at thecellvision.org/endocytosis.

#### Functional similarity of penetrance mutants and SPMs vs non-morphology-mutants

We calculated the Pearson’s correlation for every pair of phenotype profiles as described above. Next, we grouped gene pairs by different functional criteria: i) gene pairs that encoded members of the same protein complex or members of different protein complexes; ii) gene pairs that encoded proteins in the same pathway or in different pathways; iii) gene pairs that had significantly correlated genetic interaction profiles or not^13, 85^ (PCC > 0.2, GI PCC dataset downloaded from thecellmap.org); and iv) gene pairs that were co-expressed or not. Functionally related gene pairs were defined as those that belong to the same protein complex or pathway, have a significant GI profile similarity, or are co-expressed. We used one-sided Mann-Whitney U tests to evaluate whether differences between the correlation sets were significant.

#### Mean specific phenotype fraction per protein complex and within-complex PCC

For each protein complex and mutant phenotype, we calculated the mean specific phenotype fraction and standard deviation across genes encoding members of the complex. For genes with more than one allele screened, we used the mean phenotype fraction across alleles. We calculated the Pearson’s correlation for every pair of mutant phenotype profiles for genes coding for members of the complex, and calculated the mean PCC of all complex gene pairs. Results are included in Supplementary Table 7.

#### Assessing mutant phenotype relatedness

##### Common SPMs between mutant phenotypes

For each pair of phenotypes, we evaluated whether they tended to share more stringent SPMs than expected by chance. We calculated p-values using one-sided Fisher’s exact tests, and the False Discovery Rate (FDR) to correct for multiple tests. Additionally, for every phenotype we built a profile using specific phenotype fraction values of all genes. Next, we computed Pearson’s and Spearman’s correlation across all pairs of phenotype profiles. Results are included in Supplementary Table 5.

##### Enrichment of protein complexes between mutant phenotypes

For each pair of phenotypes, we retrieved the set of SPMs that shared both phenotypes and calculated if the common set was enriched for protein complexes. Protein complexes with p-value < 0.01 and at least 2 shared protein complex components were considered significant. Additionally, we required the overlap of common SPMs with members of a complex to be higher than 95% of the overlaps obtained in randomized phenotype-gene networks. Results are included in Supplementary Table 5.

### Quantification of follow-up experiments related to the assessment of incomplete penetrance

#### Quantifying penetrance as a function of replicative age

CellProfiler was used for cell segmentation and quantitative feature extraction (including cell size and mean WGA intensity). A 2NN was used to assign each cell a phenotype class and WGA intensity was used as a proxy for replicative age. For each mutant strain, cells were sorted based on their mean WGA intensity, and grouped into 6 bins as follows: 50% of cells with the lowest mean WGA intensity corresponding approximately to virgin daughters; the next 25% of cells corresponding approximately to mother cells that had undergone 1 division; the next 12.5% of cells corresponding approximately to mother cells that had undergone 2 divisions; and so on, up to the last bin containing 3.13% of the cell population with the highest WGA intensities that were assumed to have undergone 5 or more cell divisions. For each strain and aging bin we then determined the fraction of outliers. On average, approximately 3800 cells were analysed for each strain for each replicate.

#### Effect of the UPR pathway: Clustering of penetrance profiles

CellProfiler was used for cell segmentation and quantitative feature extraction (including mean UPRE-mCherry intensity). OC-SVM outlier detection was used to assign each cell to the wild-type or outlier group. For each mutant strain, cells were sorted based on their stress response-level (mean UPRE-mCherry signal intensity), and grouped into 10 bins of equal cell numbers. For each bin we determined the fraction of outliers (unsupervised penetrance). STEM software^103^ was used to cluster mutant strains into groups with distinct UPR profiles using k-means clustering. On average, 600 cells were analysed for each strain.

### Data and software availability

#### Data Resources

All penetrance and phenotype results are available at: https://thecellvision.org/endocytosis.

Normalized feature data of single cells used for neural network training, and additional files that support the analyses are available at: https://thecellvision.org/endocytosis/supplemental.

All raw extracted quantitative features of segmented single cells will be provided upon request.

#### Images

All images are available for browsing at: https://thecellvision.org/endocytosis. Batch transfer of raw images will be provided upon request.

#### Source code

Code for the single cell labelling tool, unsupervised ocSVM for outlier detection, and 2 hidden-layer fully connected neural network for single-cell classification is available at: https://thecellvision.org/tools.

Further information and requests for resources and reagents should be directed to and will be fulfilled by Brenda J. Andrews.

## ACKNOWLEDGMENTS

We thank Michael Costanzo, Jing Hou, Sheena C. Li, Bryan-Joseph San Luis, Jolanda van Leeuwen, and Chad Myers for technical help and critical comments. This work was primarily supported by grants from the Canadian Institutes for Health Research (FDN-143264 and -143265 to BA and CB respectively) and the National Institutes of Health (R01HG00583). CB holds a Canada Research Chair (Tier 1) in Proteomics, Bioinformatics & Functional Genomics. PA acknowledges financial support from the Spanish Ministerio de Economía y Competitividad (BIO2016-77038-R) and the European Research Council (SysPharmAD: 614944). CP is supported by a Ramon y Cajal fellowship (RYC-2017-22959). QM holds a Canada CIFAR AI chair at the Vector Institute and is supported by an Ontario Institute for Cancer Research Investigator award.

## AUTHOR CONTRIBUTIONS

M.M.U, H.F. and E.S. carried out and analysed experiments. M.M.U, N.S. and C.P. performed large-scale data analysis and interpretation. M.M.U, N.S., M.U., C.P., P.A. and A.S. performed additional data analysis. N.S., M.U. and M.P.M. developed the computer code, software and databases. M.M.U, N.S., H.F. and C.P. created figures. M.M.U, H.F., N.S., Q.M., C.B. and B.J.A. wrote the manuscript. Q.M., C.B. and B.J.A. supervised the project. M.M.U, C.B. and B.J.A. designed the project.

## SUPPLEMENTARY FIGURES

**Supplementary Figure 1:**
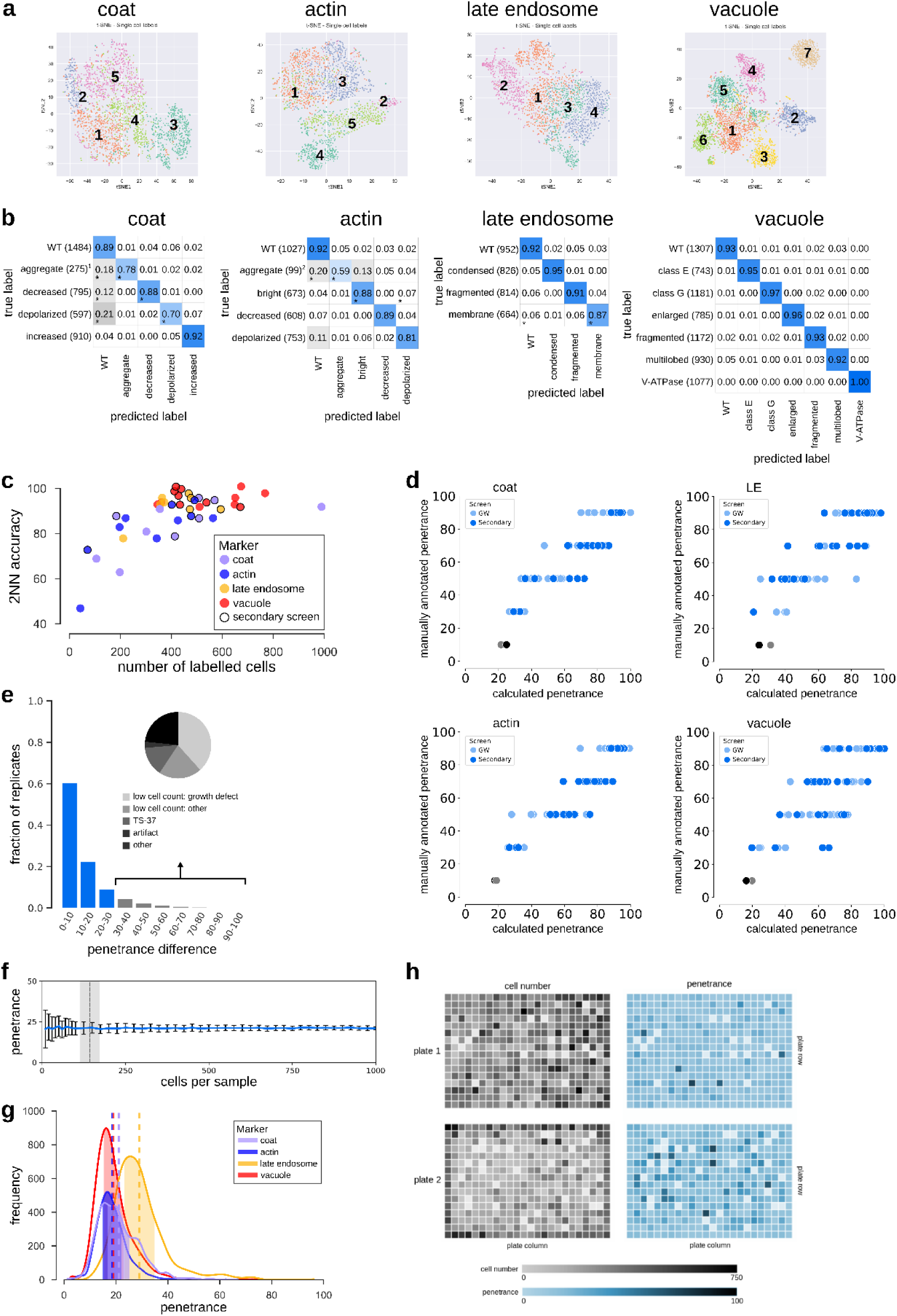
Factors affecting classification accuracy and penetrance. Related to Figure 1, Supplementary Table 1. (a) High-dimensional feature space for single cells (color-coded by phenotype) from the training sets visualized with 2D t-SNE. Numbers follow the phenotype order listed in Figure 1c. (b) Confusion matrices illustrating the classification accuracies of the 2NN classifiers for all phenotypes. Numbers in the matrix reflect the mean accuracy of both genome-wide and secondary screens. * denotes phenotypes where the difference in accuracy between the genome-wide and secondary screens was greater than 0.10. Numbers in brackets indicate the total number of labelled cells in the two filtered training sets for each phenotype. The classifiers for the two ‘aggregate’ phenotypes (denoted^1, 2^) were trained using less than 100 labelled cells in one or both of the screen types. The intensity of the blue colour in each block of the matrix indicates the fraction of cells classified from each class predicted to be in a given class (scale bar to the right). Classification accuracy for each class is indicated by the number in each block. (c) Scatter plot showing the 2NN classifier accuracy and number of labelled cells for each training set separately (N = 42), where each dot represents one phenotype class. No outline: training set for genome-wide screen. Black outline: training set for secondary screen. (d) Comparison of the manually assigned and computationally derived penetrance of positive control strains (see Supplementary Table 1 for list of strains). Each dot represents one positive control from either the genome-wide (GW) screens (light blue dots) or secondary screens (dark blue dots), and grey dots are wild-type controls. LE = late endosome. (e) Analysis of penetrance in biological replicates. The bar graph shows the fraction of biological replicates grouped according to their difference in penetrance (N = 15398 replicate pairs). Less than 10% of replicates have a penetrance difference > 30 (grey bars), with an average penetrance difference of 11.2. Insert pie chart shows a break-down of the underlying cause of large penetrance differences. (f) Bootstrapping on wild-type cell populations to determine the number of cells sufficient to obtain a confident penetrance calculation. The shaded area indicates the range of the minimum sample size across the four screened markers (defined as the sample size where the relative standard deviation falls below 0.2). Data are presented as the mean penetrance across 100 independent samplings for each sample size (blue line) +/-SD (error bars). (g) Penetrance frequency distribution of wild-type replicates for each of the four markers extracted from genome-wide screening data. The shaded area indicates the mean (vertical dashed lines) +/-0.2 x mean. Colours represent the different endocytosis markers as shown in the legend. (h) Evaluation of possible batch effects in the penetrance analysis. Representations of two screened plates illustrating cell count (grey) and computationally derived penetrance (blue) in each well are shown. A darker shade (of grey or blue) indicates increased cell number or penetrance as shown on the key below the plate representations. Even though uneven growth conditions can lead to plate-layout effects, such as gradients (top plate) or more favourable edge conditions (bottom plate), the cell density differences due to experimental artifacts do not significantly affect penetrance analysis.

**Supplementary Figure 2:**
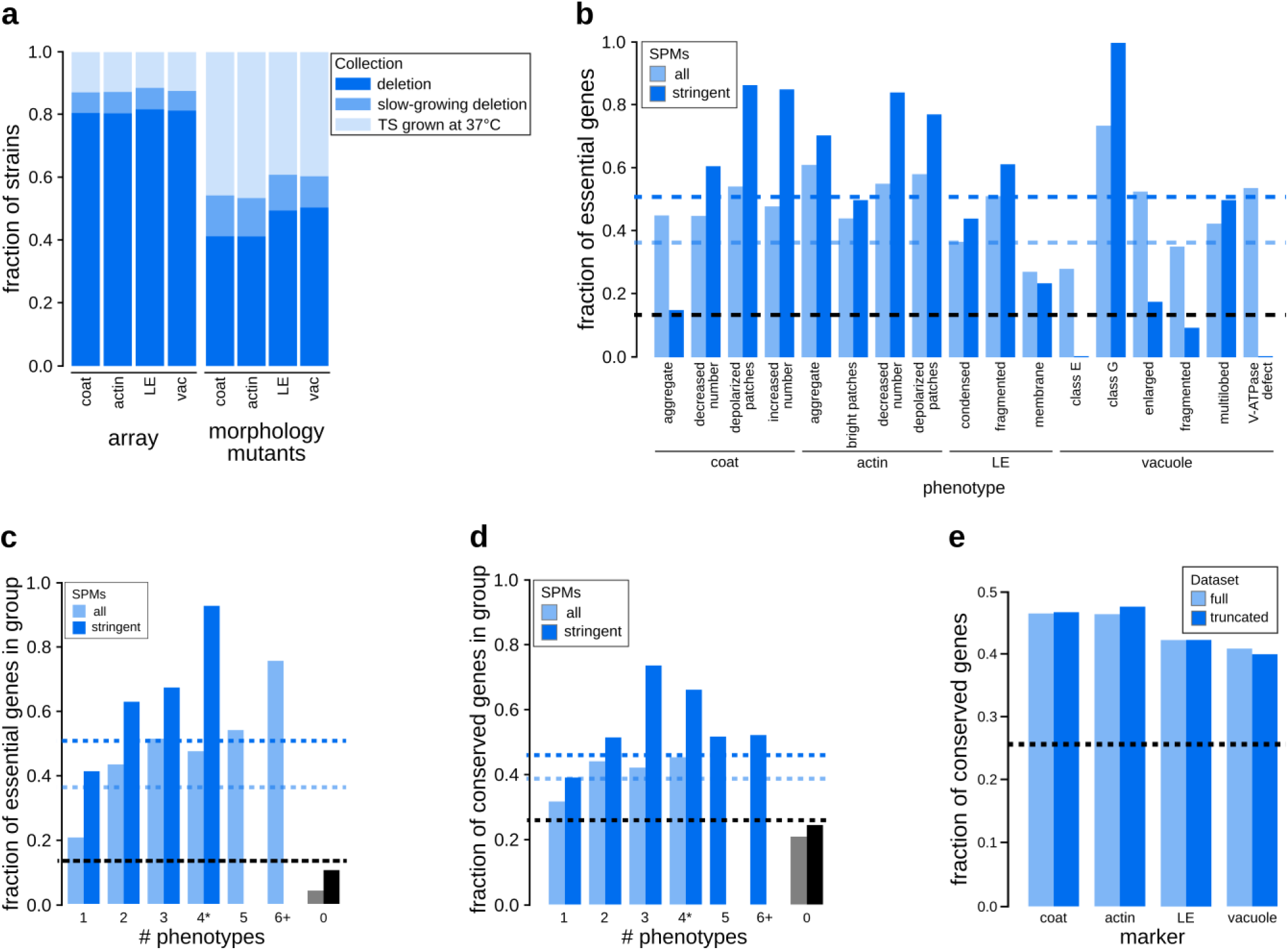
Emerging properties of mutant phenotypes. Related to Figure 2, Supplementary Table 2. (a) Comparison of the fraction of mutants screened and the fraction identified as morphology mutants in each strain collection. Stacked bar graphs show the fractions of strains in the screened array (array) and in the set of morphology mutants grouped based on the mutant strain collection for each individual marker (deletion mutant collection – dark blue; slow-growing nonessential gene deletion collection – medium blue; collection of strains with TS mutations in essential genes – light blue). LE: late endosome; vac: vacuole. (b) Relationship between specific phenotype mutants (SPMs) and essential genes. Bar graph showing the fraction of essential genes in sets of SPMs (light blue) and stringent SPMs (dark blue) for each individual phenotype. The black dashed line represents the fraction of essential genes in the screened mutant array. Blue dashed lines indicate the fraction of essential genes for all SPMs (light blue) and stringent SPMs (dark blue). LE: late endosome. (c) Bar graph illustrating the relationship between essential genes and morphological pleiotropy. Bar graph showing the fraction of essential genes in specific phenotype mutants (SPMs; light blue) and stringent SPMs (dark blue) grouped by the number of phenotypes they affect. Blue dashed lines indicate the fraction of essential genes for all SPMs (light blue) and stringent SPMs (dark blue). The black dashed line represents the fraction of essential genes in the screened mutant array. (d) Relationship between conserved genes and morphological phenotypes. Bar graph showing the fraction of conserved genes in specific phenotype mutants (SPMs; light blue) and stringent SPMs (dark blue) grouped by the number of phenotypes they affect. The black dashed line represents the fraction of conserved genes in the screened mutant array. Blue dashed lines indicate the fraction of conserved genes for all SPMs (light blue) and stringent SPMs (dark blue). (e) Bar graph showing the fraction of conserved genes in our morphology mutant sets for each of the markers for the full dataset, and a truncated dataset with excluded genes annotated to GO Slim biological process terms associated with endocytosis and the endomembrane system. Black dashed line denotes the fraction of conserved genes in the screened mutant array. LE: late endosome.

**Supplementary Figure 3:**
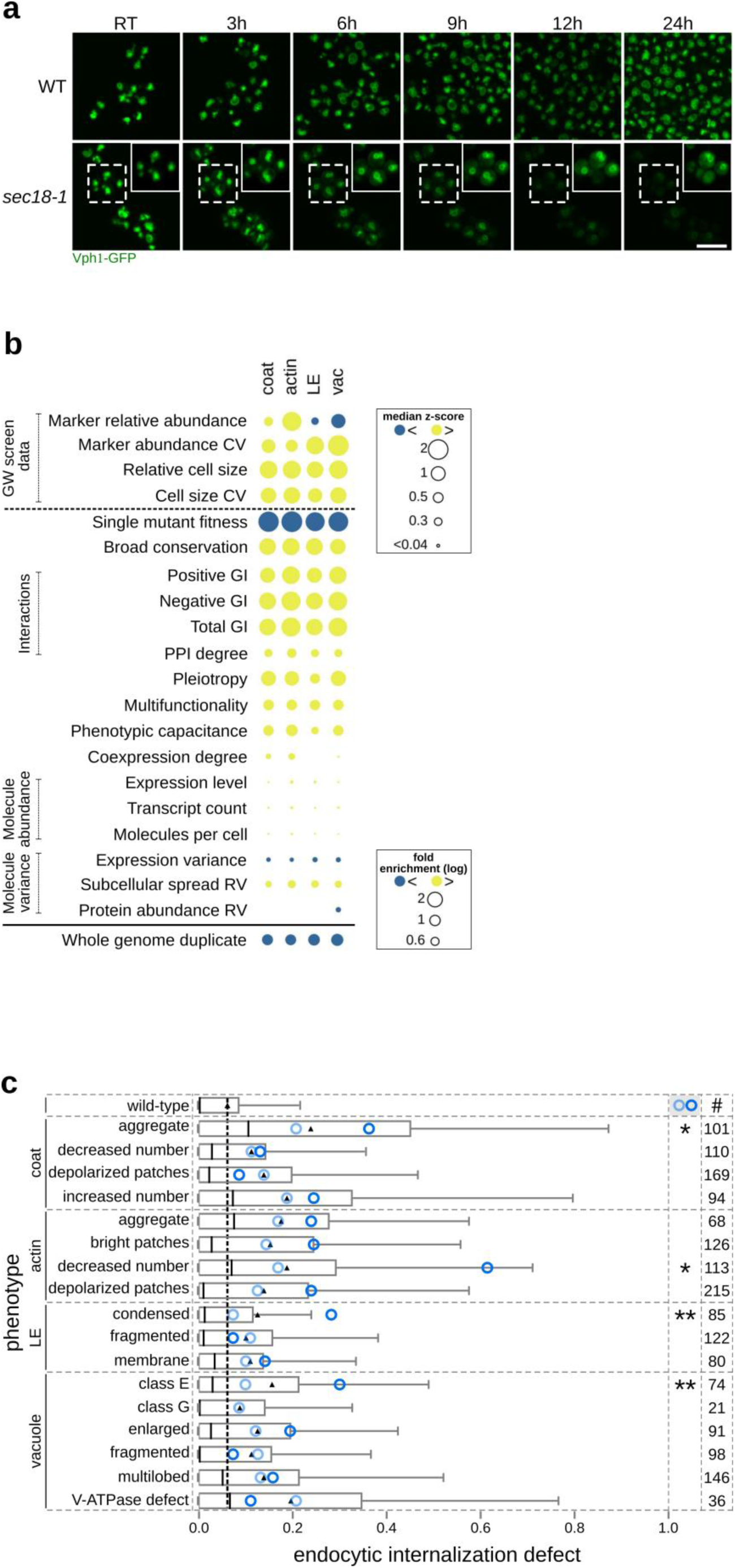
Properties of specific mutant phenotypes. Related to Figure 3, Supplementary Table 3, Supplementary Table 6. (a) Time-course analysis of vacuolar class G phenotype formation. Wild-type and *sec18-1* strains expressing Vph1-EGFP were first imaged at room temperature (RT), the temperature was then shifted to 37°C, and images were acquired at the indicated time points (in hours after shift). Signal intensity of the magnified inserts (in solid boxes within the micrographs) was adjusted to optimize phenotype visualization. Scale bar: 10 µm. (b) Gene feature enrichment analysis of the morphology mutants for each endocytic marker. Significance was determined using one-sided Mann-Whitney U tests for numeric features, and one-sided Fisher’s exact tests for binary features. For numeric features, dots represent median z-score normalized values. For binary features (below the solid black line), dots represent fold enrichment. Gene features derived from our genome-wide screens are indicated with “GW screen data” (shown above the black dotted line). CV: coefficient of variation. GI: genetic interaction. RV: relative variability. LE: late endosome; vac: vacuole. (c) Horizontal bar graph showing the distribution of endocytic internalization defect (invertase score as assessed in (Burston et al., 2009)) for nonessential specific phenotype mutants (SPMs). Several phenotypes show a significant difference between SPMs with a high specific phenotype fraction (dark blue circle) compared to those with a lower specific phenotype fraction (light blue circle). *, ** denote phenotypes with a significant difference between the two groups (p-value < 0.05, or < 0.01; significance was calculated using Kolmogorov-Smirnov tests). Black triangle: mean; black line: median; black dashed line: mean of phenotypically wild-type mutants. Numbers in the right-most column indicate the number of genes included in the analysis. Whiskers extend to the 5^th^ and 95^th^ percentile. LE: late endosome.

**Supplementary Figure 4:**
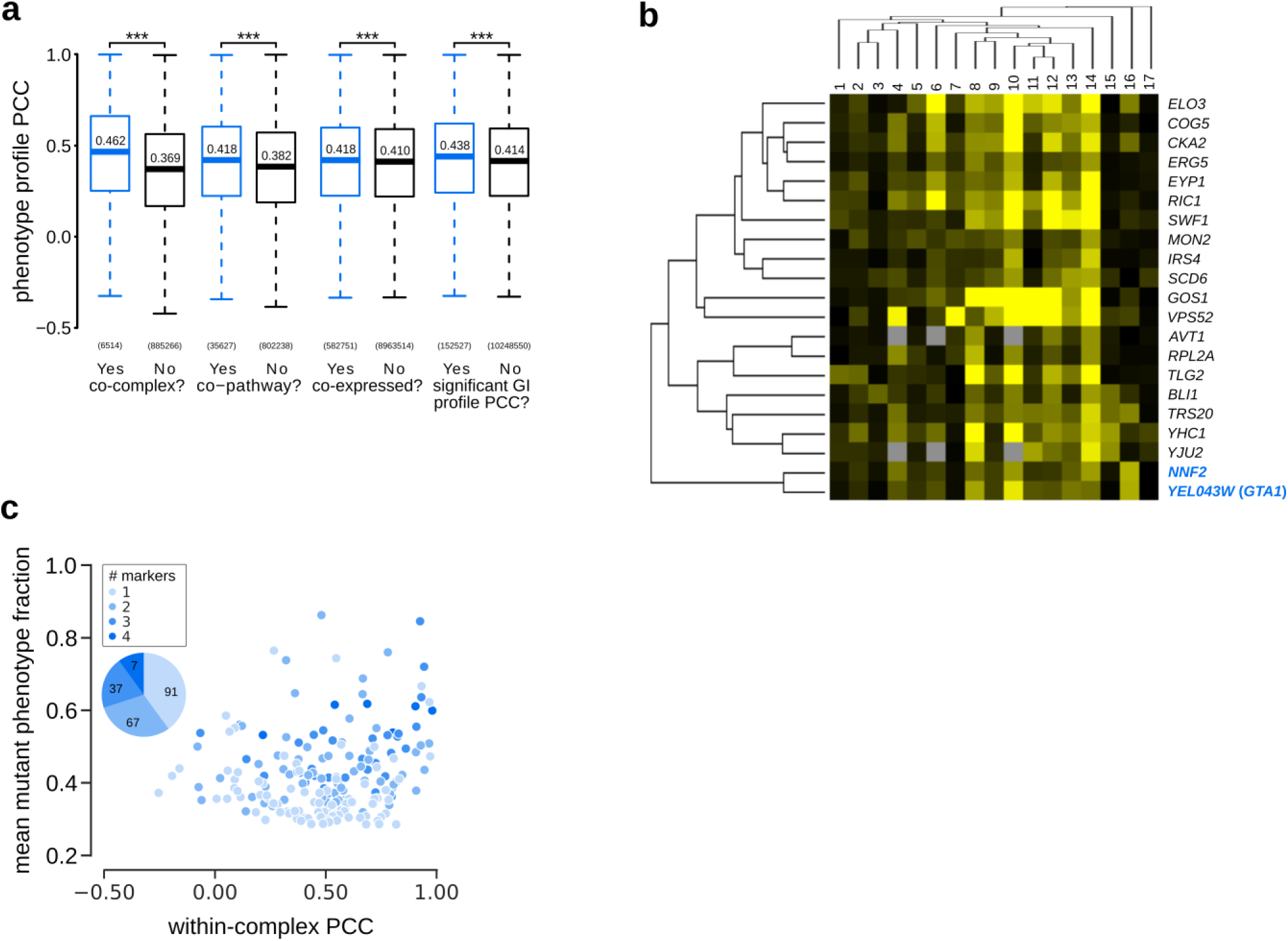
Relationship between phenotype profiles and functionally related gene pairs. Related to Figure 4, Supplementary Table 7. (a) Phenotype profile similarity of functionally related pairs of genes. Box plot indicates the distribution of Pearson correlation coefficients (PCCs) between pairs of specific phenotype profiles for genes encoding members of the same or different protein complex (co-complex); proteins in the same or different pathway (co-pathway); genes that are co-expressed or not (co-expressed), and gene pairs that have a significant GI profile similarity or not (significant GI profile PCC). The number of pairs evaluated in each set is shown on the x-axis. Significance was determined using one-sided Mann-Whitney U tests. ***p-value < 0.001. (b) Phenotype profile cluster containing *NNF2* and *YER043W* (*GTA1*) (highlighted in blue). Phenotypes 1-17: [1] coat: increased patch number; [2] coat: aggregate; [3] vacuole: class E; [4] late endosome: condensed; [5] actin: bright patches; [6] late endosome: membrane; [7] actin: aggregate; [8] coat: decreased patch number; [9] actin: decreased patch number; [10] late endosome: fragmented; [11] coat: depolarized patches; [12] actin: depolarized patches; [13] vacuole: multilobed; [14] vacuole: fragmented; [15] vacuole: enlarged; [16] vacuole: class G; [17] vacuole: V-ATPase defect. (c) Relationship between protein complexes and morphological phenotype profile correlations. Scatter plot showing mean mutant phenotype fraction (Y axis) and mean within-complex phenotype profile PCCs (Pearson Correlation Coefficient; X axis) for individual protein complexes (n = 202). The insert pie chart shows the proportion (and number) of protein complexes that affect 1, 2, 3, or all 4 markers. Mean penetrance was calculated only from affected markers. Complexes are color-coded based on the number of markers they affect.

## SUPPLEMENTARY TABLES

**Supplementary Table 1:** List of positive controls for each marker and screen type, information on training set size and 2NN accuracy. Related to Figure 1 and Supplementary Figure 1.

**Supplementary Table 2:** Main results table. Contains data on phenotype, penetrance and threshold information for each screened mutant strain, and consensus morphology mutant lists. Related to Figure 1, Figure 2, Figure 3, and Figure 5.

**Supplementary Table 3:** Enrichment results of the morphology mutants. Related to Figure 2, and Supplementary Figure 3.

**Supplementary Table 4:** Protein complex, localization, cell size and marker abundance standards. Related to Figure 2, Figure 3, Supplementary Figure 3, and Supplementary Figure 4.

**Supplementary Table 5:** Analysis of the common morphology mutants of endocytic compartment phenotypes. Related to Figure 3.

**Supplementary Table 6:** Endocytic internalization defect comparison of the mutant phenotypes. Related to Supplementary Figure 3.

**Supplementary Table 7:** Within-complex phenotype profile similarity. Related to Supplementary Figure 4.

**Supplementary Table 8:** Factors contributing to incomplete penetrance. Related to Figure 5, Figure 6.

Supplementary tables 1-8 are available at https://thecellvision.org/endocytosis/supplemental.

**Supplementary Table 9:**
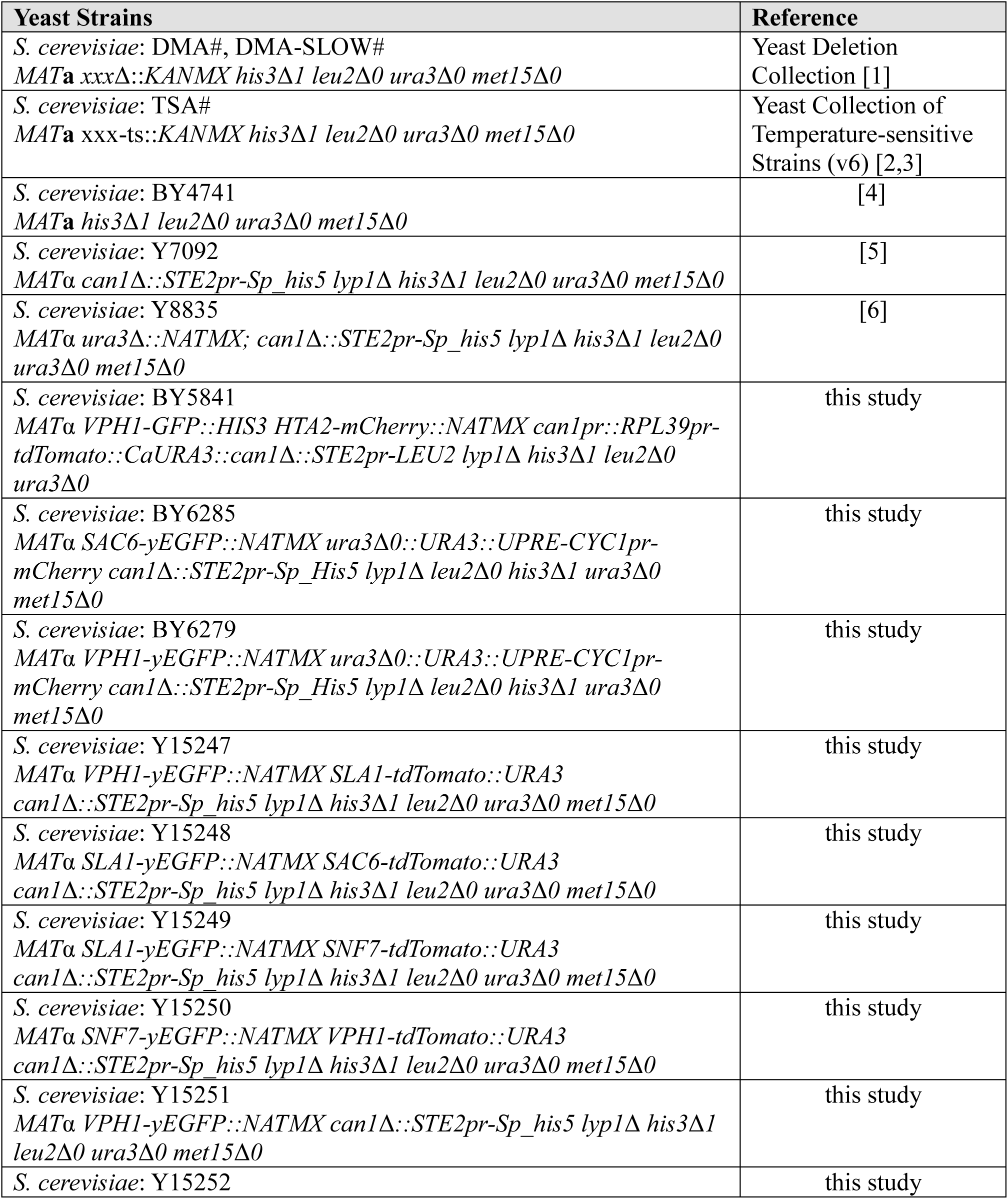
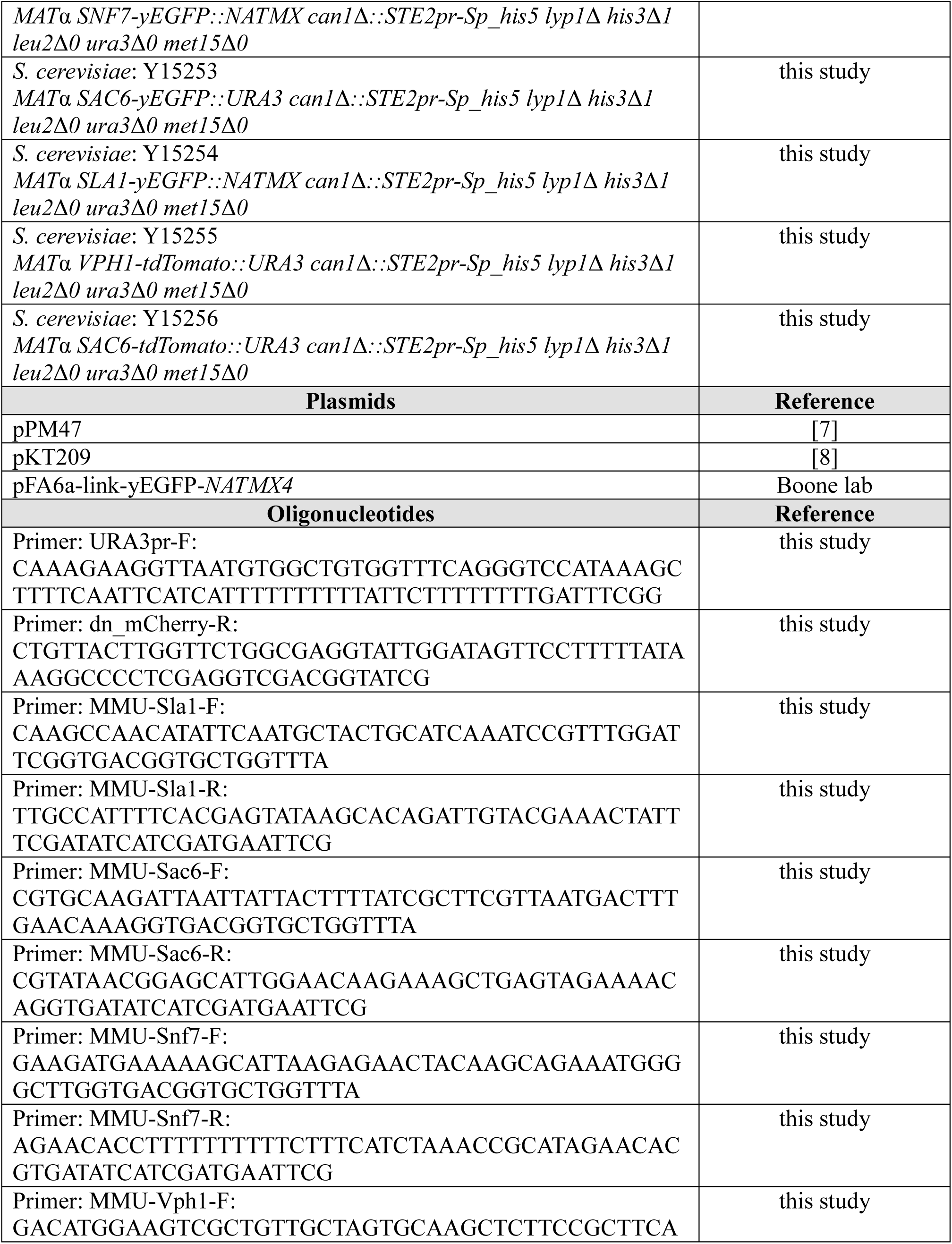
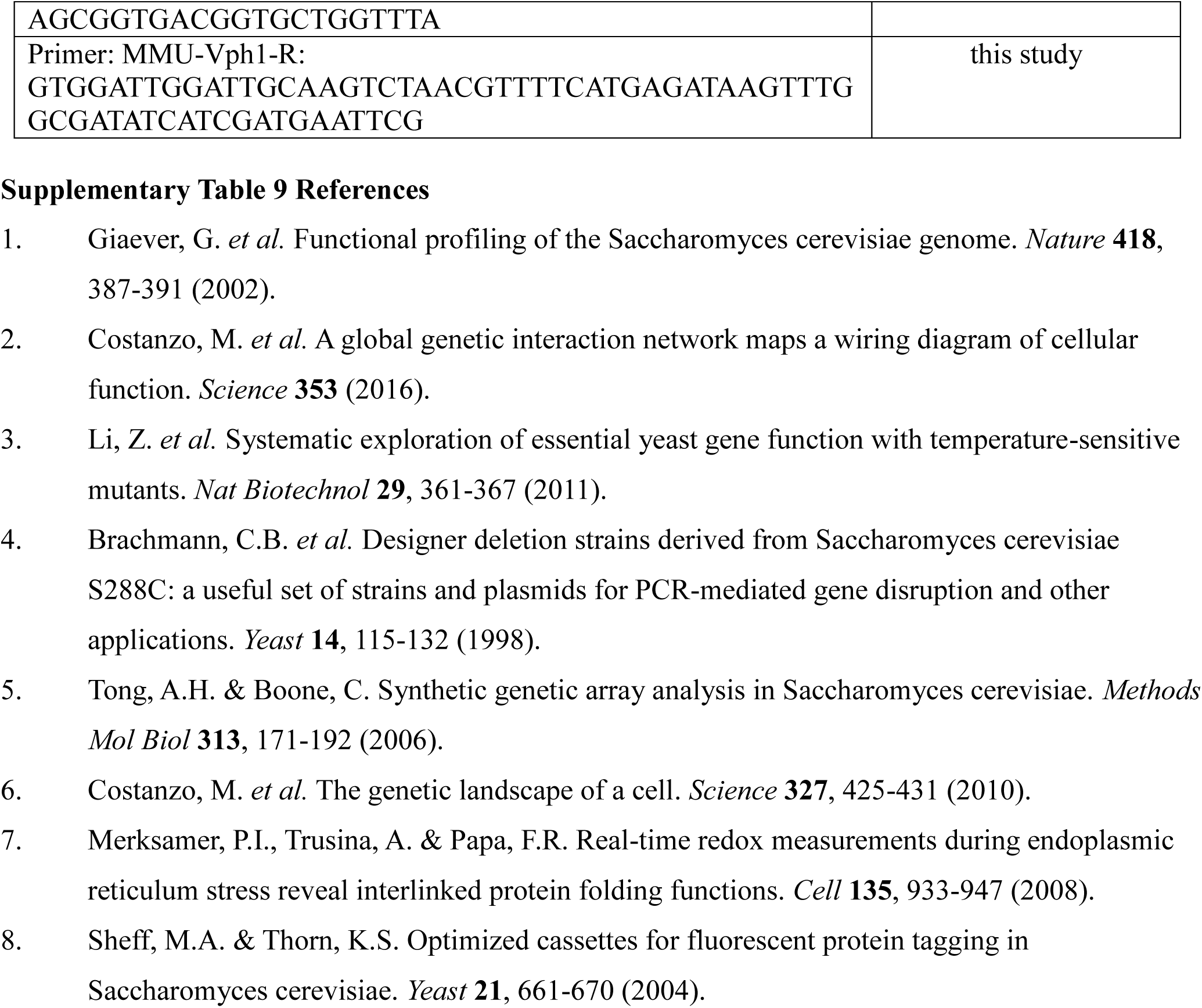
List of strains, plasmids and oligonucleotides used in this study.

